# A structural roadmap for the formation of the coronavirus nsp3/nsp4 double membrane vesicle pore and its implications for polyprotein processing and replication/transcription

**DOI:** 10.1101/2025.04.10.648184

**Authors:** Jason K. Perry, Samuel Itskanov, John P. Bilello, Eric B. Lansdon

## Abstract

Coronavirus replication is understood to occur within double membrane vesicles (DMVs) that arise during viral infection. Prior work has determined that these DMVs have characteristic pores formed from the non-structural viral proteins nsp3 and nsp4, which facilitate export of newly synthesized viral RNA. Yet how the replication machinery, which is comprised of the non-structural proteins nsp7 to nsp16, is recruited to the DMV remains a mystery. Working from AlphaFold and previously determined structures, we constructed a series of models that link formation of the DMV pore to nsp5 protease processing of the polyprotein and trapping of the cleaved products within the DMV itself. We argue that the initial pore is formed from twelve subunits of nsp3 and six subunits each of the intermediate uncleaved polyproteins pp1a’ (nsp4-nsp10) and pp1ab’ (nsp4-nsp16). Formation of this initial structure activates the protease function of alternating nsp5 subunits within a close-packed ring, facilitating the initial *trans-* cleavage of the nsp4-nsp5 linkage. Maturation of the pore follows, as does formation of canonical nsp5 dimers, which can process the remainder of the polyproteins. When protease activation occurs subsequent to closure of the DMV, the cleavage products, whose stoichiometry is consistent with the previously proposed nsp15-centered hexameric replication complex, will be trapped inside.

**Importance:** Coronaviruses, like many other positive-sense single-stranded RNA viruses, rely on intracellular membrane remodeling to create a protected environment for efficient replication to occur. The resulting double membrane vesicles (DMVs) have characteristic pores formed from the non-structural viral proteins nsp3 and nsp4, while enzymes responsible for RNA replication, including the polymerase nsp12, are contained inside. Recent structural work has elucidated the nature of the pores, but how the polymerase and other viral proteins are recruited to the DMV represents a major gap in our current knowledge. Here we present a novel, step-by-step structural model of how the pores initially form from the uncleaved polyprotein, how protease activity is initiated, and how the viral replication machinery is trapped within the DMV.

## Introduction

A unifying feature of the cytosolic replication of single-stranded (+) RNA viruses is a remodeling of host intracellular membranes to create local protective environments which facilitate the generation and storage of nascent viral RNA.(1) Chief among these modifications is the creation of replication organelles (ROs) which can be categorized as either invaginated spherules, as in the case of flaviviruses, or double membrane vesicles (DMVs), as in the case of enteroviruses.

Coronaviruses are an example of the latter, where remodeling of the endoplasmic reticulum (ER) membrane produces a network of DMVs. Accumulating evidence points to these vesicles as housing both viral RNA and the proteins necessary for replication,(2–8) thus serving the dual purposes of shielding the RNA from triggering an immune response and protecting the RNA and viral proteins from degradation. Research has focused on the viral transmembrane non-structural proteins nsp3, nsp4 and nsp6 as the principal drivers of membrane remodeling and DMV formation.(9) The general picture that has emerged from this work is that nsp3 and nsp4 are primarily responsible for membrane doubling and DMV formation, while nsp6 is responsible for membrane zippering and the tethering of DMV clusters to lipid droplets and the ER.(10) Initial images of severe acute respiratory syndrome (SARS) DMVs suggested a closed organelle,(11) however more detailed cryo-electron tomography (cryo-ET) images have captured multiple distinct pores embedded in the membranes of each DMV, providing a necessary mode of egress for newly synthesized RNA. Low resolution cryo-ET reconstructions for both murine hepatitis virus (MHV)(12) and SARS-CoV-2(13) clearly revealed pore structures with distinct six-fold symmetric cytosolic crowns terminating in prominent prong-like features. In the case of SARS-CoV-2, this cryo-ET work also demonstrated that DMV formation and pore formation go hand in hand, and that nsp3 and nsp4 are the minimal viral components required. A recent 4.2 Å resolution cryo-ET structure further revealed the pore in even greater detail, specifically showing the crown to be composed of twelve subunits of nsp3, anchored to a dodecameric arrangement of nsp4 spanning both inner and outer membranes.(14).

Nsp3 and nsp4 can be found on the first open reading frame (ORF1) of the coronavirus genome. Encompassing over 70% of the genome, ORF1 encodes two viral polyproteins, pp1a (nsp1-nsp10) and pp1ab (nsp1-nsp16). They are distinguished by a ribosomal frameshift located between nsp10 and nsp1*2*.(15) Both polyproteins are cleaved by viral proteases embedded in nsp3 and nsp5. Studies of MHV and SARS established that the first three non-structural proteins (nsp1-nsp3) are efficiently cleaved by the papain-like proteases of nsp3 (PLpro in SARS-CoV-2),(16, 17) while the remainder of the polyprotein (nsp4-nsp10, referred to here as pp1a’, and nsp4-nsp16, referred to as pp1ab’) are cleaved in a more regulated manner by the main protease (Mpro) of nsp5.(18, 19) Notably, biochemical studies of cleaved nsp5 have shown the protease to be significantly more active when dimerized,(20) but the factors that initiate nsp5 maturation and polyprotein processing have yet to be elucidated.(21) Considering that uncleaved nsp5 is sandwiched between the membrane binding proteins nsp4 and nsp6, additional constraints on its initial activity should be expected, but these constraints have not been explored and remain poorly understood. With respect to the DMV pore, it has been clearly demonstrated that cleavage between nsp3 and nsp4 is required prior to pore formation,(13, 22) but no comparable study has established when in the process of pore formation cleavage between nsp4 and nsp5 occurs. Thus, whether nsp5 maturation occurs prior to pore formation or in concert with it is an open question.

Once the pp1a’ and pp1ab’ polyproteins are fully cleaved, the resulting individual proteins, nsp7-nsp16, constitute the viral replication machinery.(23) At a minimum, RNA polymerization is carried out by a complex of (nsp12)(nsp7)(nsp8)_2_.(24) Additional enzymatic domains of nsp12,(25) nsp13,(26, 27) nsp14(28, 29) and nsp16,(30) aided by the cofactors nsp9 and nsp10, are responsible for other critical viral RNA replication functions including proofreading and mRNA capping. Previously, we proposed these viral proteins assemble around the endonuclease nsp15 to form a hexameric complex of over 60 subunits.(31) Our modeling suggested this complex efficiently coordinates the various functions just described. Implicitly, these proteins must be recruited to the DMV prior to its closure for replication inside the organelle to occur, although how that might be achieved is unknown, representing a major gap in our knowledge of coronavirus replication.(6–8).

Here we tackle the question of pore formation and cleavage of the polyprotein from a structural modeling approach. We first review our attempts at constructing a model of the nsp3/nsp4 pore structure, highlighting the successes and limitations of AlphaFold in such an endeavor. We ultimately propose modest changes to the modeled prongs in the published structure of Huang, et al.(14) We then examine constraints on nsp5 protease activity imposed by the flanking membrane binding proteins nsp4 and nsp6 and describe our discovery of what appears to be a polyprotein precursor to pore formation which overcomes those constraints. We propose that when this pore precursor is formed from six heterodimers of pp1a’ and pp1ab’, the cleaved products will be trapped within the DMV and have the proper stoichiometry to form the previously proposed hexameric replication complex. The implications of the proposed model are far ranging, with direct relevance to membrane remodeling, regulation of nsp5 protease activity, and formation of the replication and transcription complex within the closed vesicles.

## Results

### SARS-CoV-2 nsp3 monomer

At 1945 residues, nsp3 is the largest protein produced by the SARS-CoV-2 virus. While most of the individual domains have been solved by X-ray crystallography, AlphaFold2.1 was used to predict the full-length structure (Figure 1a). The majority of the protein, with some notable exceptions, was predicted with high confidence (Figure S1), as measured by the predicted Local Distance Difference Test (pLDDT),(32) with most domains scoring in the 70-90 range. X-ray structures for Ubl1 (ubiquitin-like domain 1)(33), Mac1 (macrodomain 1)(34), Mac3(35), DPUP (domain preceding Ubl2 and PLpro)(36), Ubl2/PLpro(37), NAB (nucleic acid binding domain)(38) and βSM (betacoronavirus specific marker)(39) were clearly reproduced in the AlphaFold model. Underscoring the success of the method, following completion of this work, a structure (PDB 8ILC) became available for the SARS-CoV-2 C-terminal CoV-Y domain (Y2, Y3 and Y4)(40) that matched the AlphaFold prediction as well. Additional domains with no available structures for any coronavirus were predicted for TM1, 3Ecto, TM2 (a bundle more accurately described as transmembrane domain 2 or TMD2) and Y1. The pLDDT score for the region between βSM and Y1, which includes TM1-3Ecto-TMD2, was comparatively lower, at 64.5, but notably the residues immediately following βSM (1318–1364) were predicted to be α-helical, with residues 1318-1349 having amphipathic character. The residues prior to TM1 (1385–1400) were predicted to form a second amphipatheic helix.

**Figure 1.**
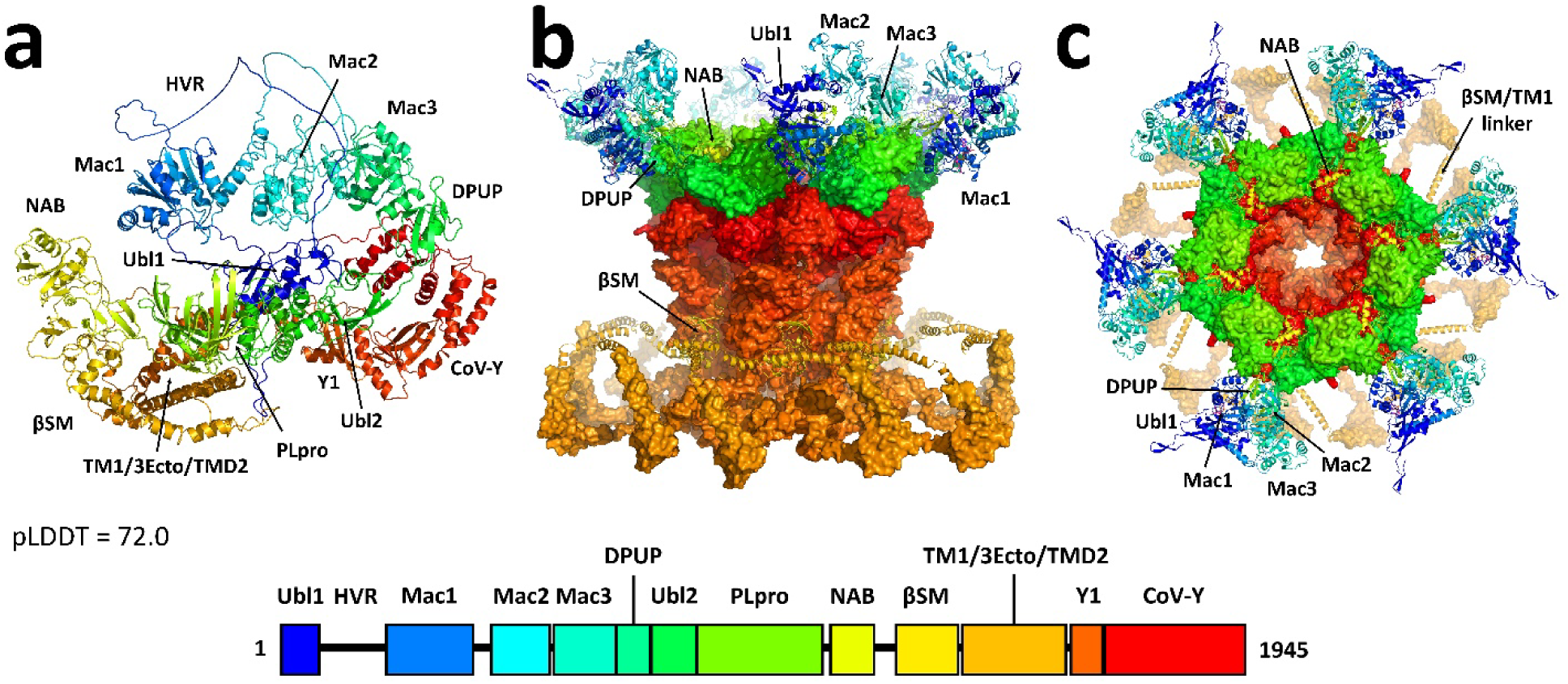
a) AlphaFold predicted structure of monomeric SARS-CoV-2 nsp3. The domain organization of nsp3 is indicated below, colored blue to red across its 1945 residues. The nsp3 structure was largely well predicted, with a few exceptions, including a long, disordered loop corresponding to the highly variable region (HVR) and a mis-folded Mac2 domain, as discussed in the text. The average pLDDT score was 72.0. b) Side and c) top views of the remodeled nsp3 dodecameric component of the DMV pore, based on the cryo-ET structure and maps of Huang, et al.(14) High resolution portions of the structure were preserved from the original fit and rendered here with surface representation. Low resolution portions of the structure were remodeled, introducing several domains that were missing from the original fit. The newly modeled domains are labeled and shown in ribbon representation here. Ubl: Ubiquitin-like domain. HVR: Highly variable region. Mac: Macrodomain. DPUP: Domain preceding Ubl2 and PLpro. PLpro: Papain-like protease. NAB: Nucleic acid binding domain. βSM: Betacoronavirus specific marker.

The most notable failure in the AlphaFold prediction was for the Mac2 domain, which deviated significantly from X-ray structures for the highly homologous SARS Mac2 domain (71% identity, 85% similarity)(41). This was unexpected, given the SARS structure’s inclusion in the AlphaFold training dataset, but it appeared to be unique to the prediction of full-length nsp3. Predictions of the isolated domain or the Mac2 and flanking domains, were consistent with the SARS structures, putting its pLDDT score in line with other domains (Figure S1). Prediction of the heterodimer of Mac2 with human Paip1 was also consistent with the existing SARS Mac2/Paip1 X-ray crystal structure (PDB 6YXJ).(42).

In general, variations in the predicted structure conformations, coupled with the pLDDT analysis, reflect significant flexibility between the domains (see Figure S2). This is most striking for the highly variable region (HVR) between Ubl1 and Mac1, which was completely disordered in the AlphaFold prediction. In addition, the loops connecting Mac1 and Mac2 (residues 388-415) and connecting NAB and βSM (residues 1197-1240) are particularly long. Ultimately, these linkers provided a set of constraints on the maximum distance and range of motion between domains that we considered when examining the nsp3 component of the pore.

### SARS-CoV-2 nsp3 dodecamer

Early analysis of the SARS-CoV-2 and MHV pore structures determined from cryo-ET suggested that the cystosolic crown is composed of a hexamer of nsp3, with the N-terminus residing in the prongs(12, 13). Due to practical limitations in the size of structures that could be predicted with AlphaFold (about 5000 residues), it wasn’t possible to generate a model for an entire nsp3 hexamer (11,670 residues) and exploration of truncated constructs had only limited success.

The publication of the 4.2 Å cryo-ET structure however provided surprising clarity.(14) The six-fold symmetric crown is instead a dodecamer of nsp3, although much of the protein structure was not resolved. Specifically, the C-terminal Y1/CoV-Y domains form the dodecameric base of the crown and the cytosolic portion of the pore. It is constructed from an inner hexameric ring of Y1/Y2 (forming the pore), which is surrounded by another hexameric outer Y1/Y2 ring to create the base of the crown. The twelve CoV-Y Y3/Y4 domains interlace above the base, forming the bulk of the crown. The crown is embedded in the outer membrane through TMD2, interacting with nsp4. Here we note that the clearest success among the AlphaFold predictions of multimeric nsp3 constructs was that of the inner Y1/Y2 hexameric ring, shown in Figure S3 in comparison to the cryo-ET structure.

The N-terminal domains preceding TM1 are only partially resolved for six subunits and not resolved at all for the other six, implying a degree of disorder, which may reflect real dynamics of the pore or the lack of other key stabilizing elements such as RNA or additional host or viral proteins. Alternatively, it may simply be an artifact of how the structure was obtained and refined (the cryo-ET structure was determined from DMVs derived from nsp3-nsp4 expression in HEK293F cells, not DMVs derived from infected cells). What is definitively resolved is that a hexameric ring of Ubl2/PLpro rests atop the CoV-Y dodecamer to build up the crown. Huang, et al. also fit each of the six prongs with the Mac2, Mac3 and DPUP domains from one subunit and the NAB domain from a neighboring subunit. However, the resolution of these prongs is considerably weaker (>9 Å), making the fit ambiguous. Several considerations led us to remodel this part of the structure. Huang, et al. assumed the prong density was primarily composed of the Mac2, Mac3 and NAB domains and used AlphaFold to predict a complex of these domains, which they docked into the density. This fit put the NAB domain implausibly far from the assumed position of the βSM domain at the base of the crown. In contrast, they did not fit the Ubl1, HVR or Mac1 domains, which Zimmermann, et al.(13) previously showed was critical to the proper formation of the prongs. Considering that Mac1 corresponds to the only Mac domain in MHV nsp3, we reasoned that it was an important component of the prong density and was likely similarly positioned in both structures., Taking into account functional constraints, similarities and differences between MHV and SARS-CoV-2, and all available cryo-ET maps, including the prior maps for both MHV and SARS-CoV-2, we fit the prongs with Ubl1, Mac1, Mac2, Mac3 and DPUP, omitting only the HVR domain for lack of structural information. Furthermore, we positioned the NAB domain between PLpro domains in the rim of the crown. We also modeled the βSM domain and two amphipathic helices on the outer membrane cytosolic surface, following previously unfit density. As a proof of principle, we connected the NAB and βSM domains with the 44-residue linker. A similar model was built for the MHV nsp3 dodecamer to establish consistency across diverse coronaviruses. The final remodeled nsp3 crown is shown in Figures 1b and 1c and described in greater detail in the Supplementary Information and Figures S4, S5 and S6.

### SARS-CoV-2 nsp4 monomer

Our model of nsp4 (Figure 2a) is consistent with previous AlphaFold models with four distinct domains.(43) The N-terminal TM1 is a single α-helix which spans residues 1-27. This is followed by the lumenal domain, a soluble region composed of two distinct lobes connected by a 9-residue linker. The previously designated TM2/TM3/TM4 helices(44) were instead predicted by AlphaFold to be a 5-helix bundle, spanning residues 277-398. For simplicity, we’ll refer to this region as transmembrane domain 2 (TMD2), with specific helices α1-α5. Finally, the AlphaFold predicted C-terminal domain (CTD) is another soluble region which reproduces existing X-ray structures(45). The average pLDDT score for the full protein was 83.0 (Figure S7).

**Figure 2.**
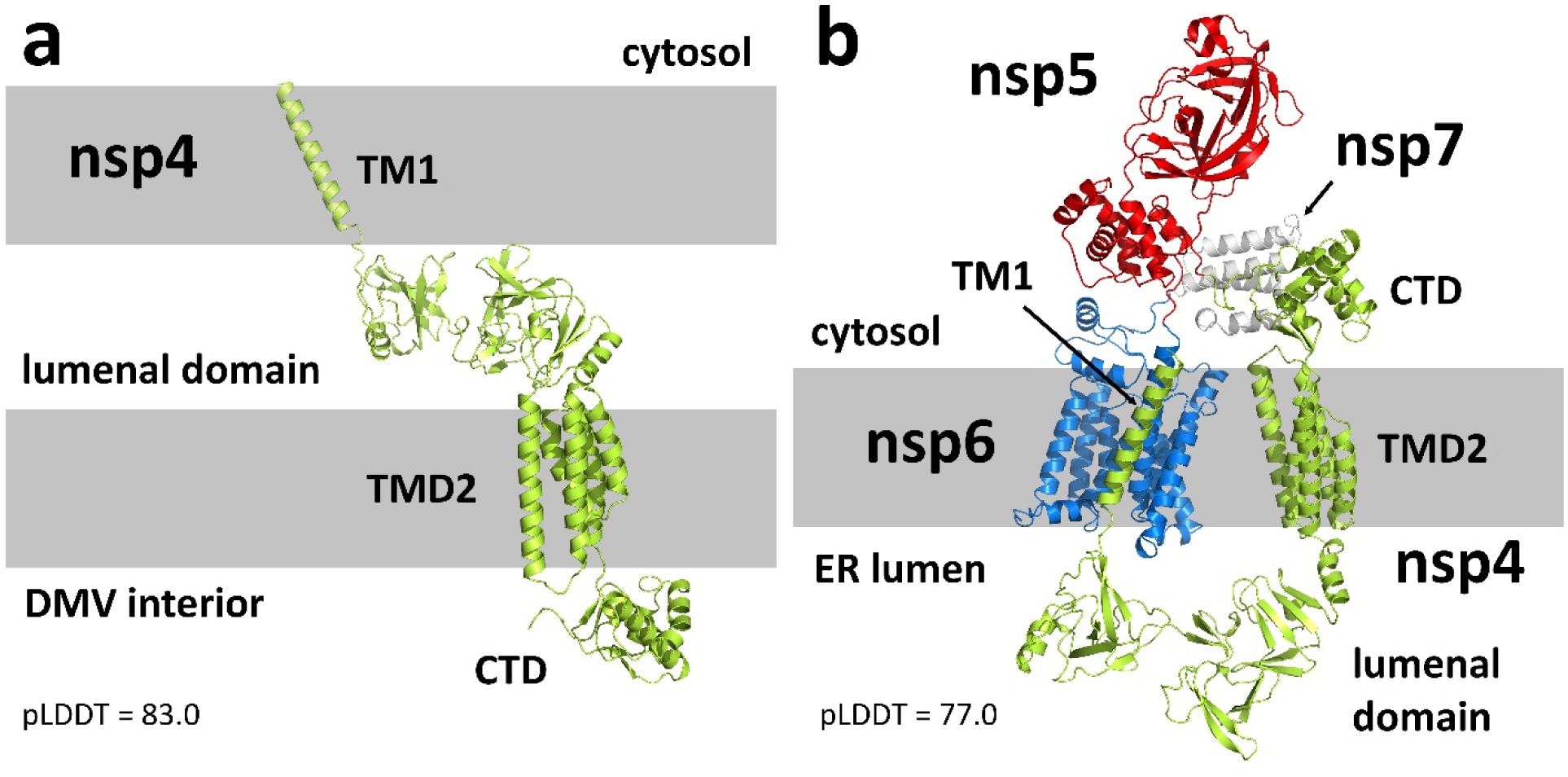
a) AlphaFold predicted structure of nsp4. The model reflects a conformation consistent with association to a double membrane (indicated by the gray bars), where the lumenal domain resides in the membrane gap. b) Predicted structure of the uncleaved nsp4-nsp7 polyprotein, with nsp4 in lime green, nsp5 in red, nsp6 in marine blue and nsp7 in white. The model adopts a conformation consistent with binding to the ER membrane, where the nsp4 N-terminus and CTD, nsp5 and nsp7 are cytosolic and the nsp4 lumenal domain is in the ER lumen. A conformational change between the N-lobe and C-lobe of the nsp4 lumenal domain determines whether the protein binds to a single membrane or a double membrane.

In none of the predicted structures does TM1 interact with TMD2. The conformation is instead consistent with nsp4 association with the DMV double membrane, where the nsp4 N-terminus is cytosolic, TM1 sits in the outer membrane, the lumenal domain is positioned in the membrane gap, TMD2 sits in the inner membrane, and the nsp4 C-terminal domain (CTD) is exposed to the DMV interior. Thus, the overall topology of the AlphaFold predicted nsp4 structure is consistent with that seen in the detailed pore structure. Details of the predicted structure, such as the lumenal domain and TMD2, were also confirmed by the cryo-ET structure, with the only notable difference being a conformational shift of helix α1 in TMD2, to be discussed below in more detail.

### SARS-CoV-2 nsp4-nsp5-nsp6-nsp7 monomer

To investigate constraints on the nsp5 protease imposed by its position within the polyprotein, we generated AlphaFold models for the uncleaved nsp4-nsp5-nsp6-nsp7 construct (Figure 2b). The pLDDT declined slightly to an average of 77.0 (Figure S8), but X-ray structures of nsp5(46) and nsp7(47) were well reproduced, and nsp6 was predicted to form an 8-helix bundle. The overall organization of the polyprotein showed some flexibility between the individual domains. However, in all predicted structures, the nsp4 TMD2 was aligned with nsp6 as if bound to a common membrane. Unlike the prediction for cleaved nsp4, the lumenal domain of nsp4 in the polyprotein adopted a conformation between its N- and C-lobes (linked by residues 124-132) which oriented TM1 such that it associated with the nsp6 helix bundle (Figure S9). This would be more consistent with association with the single ER membrane rather than the DMV double membrane, making nsp5, nsp7, the nsp4 N-terminus and the nsp4 CTD cytosolic.(48) The observation that this conformational change is driven by the nsp4 lumenal domain is consistent with previous characterizations of this domain by Klatte, et al.(49).

The nsp5 protease dimer is known to be significantly more active than the monomer,(20, 21) but a major conclusion from this work is that despite a relatively long flexible loop connecting nsp4 and nsp5 (residues 488-510), *cis*-cleavage of these proteins does not appear to be feasible. The cleavage site on the connecting loop (residues 500-501) simply cannot reach the protease active site without compromising the integrity of the protease structure (Figure 3a). Similarly, *cis*-cleavage of the nsp5 and nsp6 proteins at residues 807-808 does not appear to be possible under any circumstances. Thus, in both cases, cleavage must proceed via interaction with the protease of a second polyprotein or an already cleaved nsp5 dimer.

**Figure 3.**
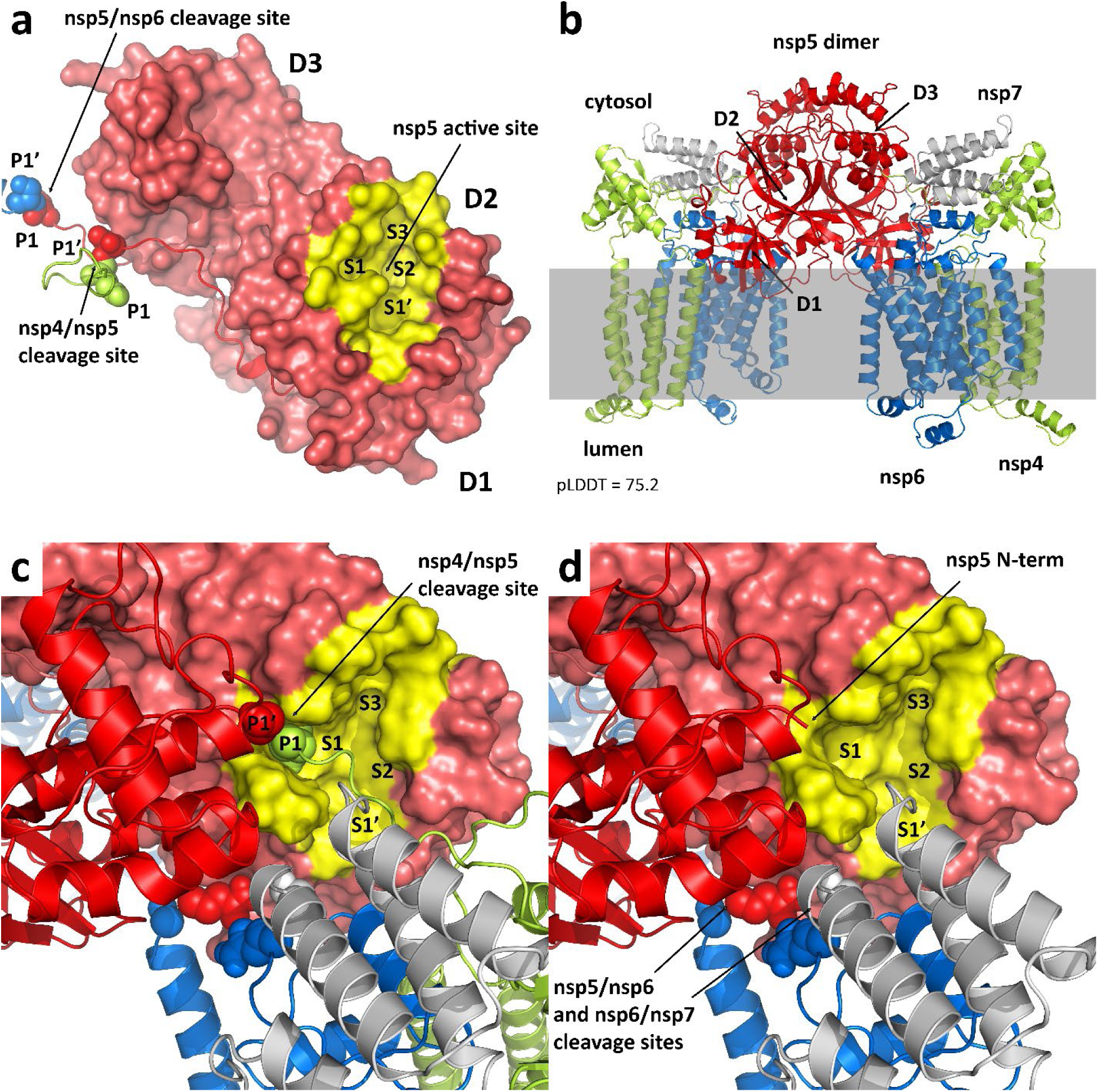
a) Location of the nsp4/nsp5 and nsp5/nsp6 cleavage sites on a single subunit relative to the nsp5 protease active site. The nsp4/nsp5 cleavage site (between residues Q500 and S501) sits on a loop that is not long enough to reasonably engage the active site. This is also true of the nsp5/nsp6 cleavage site. We can conclude that *cis*-cleavage is not a viable pathway. The three nsp5 domains are labeled D1, D2 and D3. The protease substrate recognition pockets are labeled S1’, S1, S2 and S3. The peptide cleavage sites are labeled P1’ and P1. b) AlphaFold prediction of the nsp4-nsp5-nsp6-nsp7 dimer. The predicted structure is anchored by a canonical nsp5 dimer and associated with a single membrane through nsp4 and nsp6. It has an average pLDDT score of 75.2. c) Detail of the nsp5 active site in the predicted uncleaved dimer. The nsp4/nsp5 cleavage site engages with the nsp5 active site from the other subunit, but not in a productive conformation. This effectively renders the protease inactive and implies the initial cleavage event must proceed through a different mechanism. d) Assuming nsp4/nsp5 cleavage has already occurred, the protease site becomes active, potentially capable of cleaving nsp5/nsp6 or nsp6/nsp7. The nsp5 N-terminal amine has previously been implicated in the increased activity of the dimer vs. the monomer.

### SARS-CoV-2 nsp4-nsp5-nsp6-nsp7 dimer

The cleaved nsp5 dimer has been exhaustively studied through X-ray crystallography for multiple coronaviruses.(50–52) A key factor in the increased protease activity of the dimer as compared to the monomer is how the N-terminal amine of one monomer engages with the S1 active site pocket of the other monomer. This has been shown to be critical to stabilizing the active site in both SARS(50, 53, 54) and SARS-CoV-2,(52) enhancing the protease’s catalytic activity. We considered if a canonical nsp5 dimer could form within the uncleaved nsp4-nsp5-nsp6-nsp7 polyprotein, and if so, would it be capable of self-cleavage. Using a truncated construct, in which the nsp4 TM1 and lumenal domains were removed, we generated an AlphaFold model of the uncleaved dimer possessing approximately C2 symmetry (Figure 3b). In this model, nsp4 and nsp6 are aligned as if bound to a common membrane, with the canonical nsp5 dimer oriented such that its domains 1 rest against the membrane and domains 3 are most exposed to the cytosol.

The nsp4-nsp5 cleavage site P1 Q500 residue can be seen to occupy the S1 pocket of the active site. Yet the full nsp4-nsp5 cleavage peptide is not properly aligned, and critically the P1’ S501 cleavage site residue cannot be positioned in the S1’ pocket due to constraints from the nsp5 dimer itself (Figure 3c). Thus, the dimer is incapable of cleaving nsp4/nsp5. To stress this point, a 100ns molecular dynamics (MD) simulation of the membrane bound dimer maintained this interaction between the glutamine residue and the S1 site over the entire simulation (Figure S10). We conclude from this analysis that not only does the uncleaved dimer not have the advantage of the cleaved nsp5 N-terminus to stabilize the S1 site and activate the protease it has the uncleaved nsp4 C-terminal glutamine residue blocking the site from acting on any other substrate.

If additional steric and conformational constraints from the extended polyprotein or potentially an nsp3 interaction prove to be more dominant, it should be considered that this canonical dimer may not even form until some cleavage has already occurred via a different route.

Underscoring this point, the nsp5 protein of MERS only weakly dimerizes, depending instead on a ligand-induced dimerization mechanism.(51) Indeed, as seen in Figure 3d, taking the same nsp4-nsp5-nsp6-nsp7 dimer and assuming at least one nsp4/nsp5 cleavage has occurred via some yet to be determined mechanism, the resulting nsp5 N-terminal amine then activates the protease of the uncleaved subunit. The nsp5-nsp6 and nsp6-nsp7 cleavage sites of the cleaved monomer can be observed to be in close proximity to the activated protease and could be potential substrates. This analysis suggests that an initial nsp4-nsp5 cleavage event is required but does not proceed through the canonical nsp5 dimer. Some other association between polyproteins must be responsible.

### SARS-CoV-2 nsp4 dodecamer

As with nsp3, our initial assumption was that the nsp4 component of the pore was hexameric. Based on our predicted structures for the nsp4 monomer and previous work suggesting nsp3 interacts with the nsp4 lumenal domain,(49, 55) we reasoned TM1 must sit in the outer membrane, positioning the lumenal domain in the membrane gap. This suggested the portion of the pore embedded in the inner membrane would be formed from a hexamer of the nsp4 TMD2 domains, with the nsp4 CTD exposed to the DMV interior. Thus, we worked with a truncated construct which spanned residues 256-500 of nsp4 (TMD2 and CTD). AlphaFold indeed predicted several plausible hexameric structures with pore-like properties, but none was entirely consistent with the available cryo-ET maps. However, upon close examination of the MHV map, we identified density within the membrane gap which was similar in size to the nsp4 lumen domain, but which indicated nsp4 was in fact dodecameric, not hexameric.

Unfortunately, AlphaFold predictions of dodecameric nsp4 (residues 256-500) did not produce structures with the expected C6 symmetry. The detailed pore structure from Huang, et al. confirmed that the nsp4 component is indeed a dodecamer, characterized by a central hexameric ring interlaced with an outer hexameric ring. While failing to produce the full dodecameric nsp4 assembly, AlphaFold in fact accurately predicted the structure of the inner hexameric ring, as shown in Figure S11 in comparison to the cryo-ET structure.

### SARS-CoV-2 nsp4-nsp5-nsp6 dodecamer

As the initial nsp5 cleavage events appeared to be unresolved, we considered the possibility of pore formation prior to nsp4/nsp5 cleavage. Our preliminary exploration with AlphaFold also considered hexamers of uncleaved nsp4-nsp5-nsp6 (truncated to residues 256-1042). AlphaFold generated a particularly compelling model shown in Figure 4a and Figure S12, which consisted of an inner ring of nsp4 TMD2 surrounded by an outer ring of nsp6. The two transmembrane domains are linked via an inner, close-packed solvent exposed ring formed from the nsp4 CTD, which is surrounded by a larger close-packed ring formed from nsp5. Consistent with the emerging topology of the nsp3/nsp4 pore, both nsp5 and the nsp4 CTD would be exposed to the interior of the DMV in this model.

**Figure 4.**
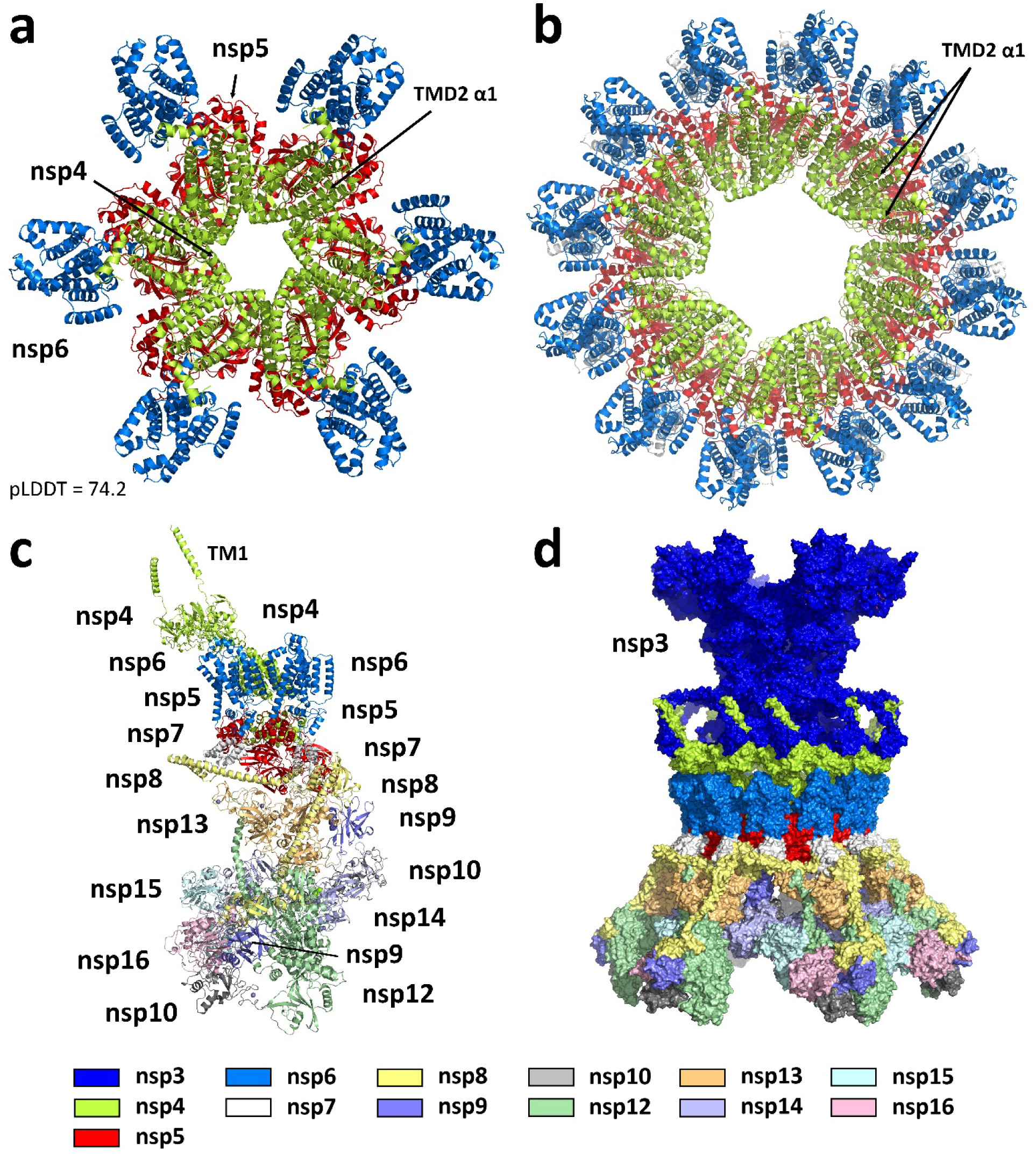
a) AlphaFold prediction of the hexameric SARS-CoV-2 nsp4-nsp6 polyprotein (a truncated construct covering residues 256-1042). Nsp4 (lime green) forms a pore-like interior, flanked by nsp6 (marine blue). Nsp5 (red) forms a close packed ring-like arrangement on the underside. b) The nsp4-nsp6 hexamer transformed into a dodecamer. The nsp4 TMD2 alternates between open and closed conformations, facilitating pore maturation upon nsp4-nsp5 cleavage. c) The pp1a’/pp1ab’ heterodimer. The dimer incorporates known interactions between the nsp proteins to the maximum extent. The nsp4 conformation is consistent with coordination to a double membrane and formation of the dodecameric ring. d) Model of the complete DMV immature pore, prior to nsp4-nsp5 cleavage. The precursor to the pore is formed from twelve subunits of nsp3 (dark blue) and six subunits each of uncleaved pp1a’ and pp1ab’.

As we came to understand the stoichiometry of the nsp4 component of the pore was dodecameric instead of hexameric, we found it was straightforward to envision similar close-packed polyprotein rings of increasing size. As an example, the chikungunya virus protein nsP1 has been shown via cryo-EM to form a dodecameric ring.(56) We used AlphaFold to look at a series of smaller aggregates (hexamer, octamer, decamer), and a similarly arranged ring was predicted in each case (see Figure S13a). Unfortunately, a dodecameric arrangement of nsp4-nsp5-nsp6 or any ring larger than a hexamer was beyond the practical limits of AlphaFold. To generate such a model, we instead resorted to a simple geometric expansion of the nsp4-nsp5-nsp6 hexamer. This was executed by defining the center of mass of each subunit within the hexamer, measuring the distance between adjacent subunits, and translating and rotating duplicate sets of these subunits to form a dodecameric ring in which this distance was maintained. Structural refinement from this starting point (as detailed below) led to the fully optimized construct shown in Figure 4b. A similar approach was applied to the AlphaFold model of hexameric chikungunya nsP1 to yield a dodecameric model which reproduces the cryo-EM structure well (Figure S13b).

In comparison to the earlier MHV and SARS-CoV-2 cryo-ET maps, this dodecameric arrangement appeared to be consistent with the overall diameter of the inner membrane pore and provided the first hint as to how these pores may form. Once the detailed pore structure of Huang, et al became available, the arrangement of the final fully cleaved pore was clear, and our attention turned to whether the uncleaved model was consistent with this structure.

As described above, the nsp4 dodecamer component of the pore structure can be described as an inner hexameric ring interlaced with an outer hexameric ring. Based on the locations of the C-terminal domains, a clear requirement for this structure to form is that the inner nsp4 hexamer must be cleaved from nsp5. However, it’s equally clear that the outer hexameric ring can remain part of the uncleaved polyprotein and retain the integrity of the pore structure. Thus, maturation of the pore within the framework of this model would at minimum require the cleavage of alternating nsp4/nsp5 sites.

This presented an interesting topological question and an opportunity to reverse engineer the process. In the nsp4-nsp5-nsp6 hexamers, we saw a domain swapping occur in which TMD2 adopts an open conformation with the α1 helix jutting out at a significant angle (∼60°) to interact with a neighboring TMD2 bundle of α2-α5. This open conformation (see Figure S14a) is also seen in both the inner and outer hexamers of the structure solved by Huang, et al., where domain swapping is clearly observed in the inner hexamer. Furthermore, examining an interlaced pair of nsp4 subunits from this structure, the open conformation has the effect of projecting one subunit inward and the other outward, causing the two to cross each other (Figure S14b).

We recognized that there was some ambiguity as to how this may manifest itself when converting the AlphaFold predicted nsp4-nsp5-nsp6 hexamer to a dodecamer. We considered three possibilities: one in which all twelve subunits adopt the open conformation, one in which they all adopt the closed conformation and one in which only alternating subunits adopt the open conformation (Figures S14c). The overall structure was unchanged except for whether the α1 helix was open or closed. From these three possibilities, it could be seen that the structure with all twelve subunits in the open conformation does not have a direct path to the interlaced final pore structure upon nsp4/nsp5 cleavage. However, in an arrangement in which only alternating subunits adopt the open conformation, upon cleavage of the nsp4-nsp5 linkages from those same six subunits, the structure would have a simple pathway to adopt the mature pore conformation (see the movie in the Supplementary Information). From this perspective, the model shown in Figure 4b may be considered a hexameric ring of polyprotein dimers, mirroring the situation for the nsp3 dodecamer.

This model was ultimately merged with the nsp4 lumenal domains of the Huang, et al. structure and our rebuilt model of the nsp3 dodecamer. The nsp4 TM1 helices, missing from the cryo-ET structure, were also added in parallel to the nsp3 TM1 helices, consistent with discernible, but previously unmodelled, density in the maps.

### SARS-CoV-2 uncleaved pp1a’/pp1ab’ model

Having created a model of dodecameric nsp4-nsp6, we considered modeling the remainder of the uncleaved polyprotein. This extended model could be derived from either pp1a’ (nsp4-nsp10) or pp1ab’ (nsp4-nsp16) or a combination of both. We considered that certain known protein-protein interactions may be preserved in this situation, creating a degree of preorganization that would facilitate efficient formation of the replication complex upon polyprotein cleavage. From cryo-EM and X-ray structures, interactions between nsp7 and nsp8, nsp7 and nsp12, nsp8 and nsp12, nsp8 and nsp13, nsp9 and nsp12, nsp10 and nsp14, and nsp10 and nsp16 are well characterized(57–60), as is nsp15 hexamerization(61).

Working from a dimer of nsp4-nsp5-nsp6 extracted from the optimized dodecameric ring, it appeared that extending both subunits to the full pp1ab’ polyprotein would be too bulky and provide only limited interactions between the subunits. In contrast, extending both to the smaller pp1a’ polyprotein would be well tolerated, but offer little in additional protein-protein interactions. However, a heterodimer formed from pp1a’ and pp1ab’ offered the maximum opportunity to incorporate the known protein-protein interactions.

Initial exploration of polyprotein flexibility was conducted with AlphaFold on multiple constructs. The uncleaved protein dimers (eg nsp9-nsp10, nsp10-nsp12, etc) revealed conformational flexibility between all pairs of proteins (Figure S15), driven by typically unstructured linkers. This led to exploration with larger fragments, with particular attention to forming interactions that are already known. We extended the initial nsp4-nsp5-nsp6 dimer to include nsp7, consistent with a conformation seen in the AlphaFold prediction of monomeric nsp4-nsp5-nsp6-nsp7. We then found we could extend one of the subunits to include nsp8 and have that protein interact with nsp7 from the other subunit just as it does in the cryo-EM structures of the replication complex (eg 6XEZ).(57) Furthermore, we found we could associate nsp13 with this nsp7/nsp8 pair, again as seen in the replication complex structure (Figure S16a). This immediately established constraints on where nsp9, nsp12, nsp14 and the second nsp8 would need to connect.

AlphaFold predictions of uncleaved nsp8-nsp9-nsp10-nsp12 led to one of the largest building blocks of the polyprotein heterodimer. The predictions showed variability in the positioning of nsp9 and nsp10, but consistency in the interaction between nsp8 and nsp12 (Figure S16b). This mirrored the replication complex as well and put constraints on how this string of proteins from pp1ab’ would be placed relative to nsp7 and nsp13. As both nsp14 and nsp16 are known to coordinate to nsp10, we further aimed to find a conformation that would allow both protein pairs to be satisfied. We proposed that one of these proteins would coordinate *cis* to nsp10 within the larger pp1ab’ polyprotein and the other would coordinate *trans* to the terminal nsp10 of pp1a’. We examined both possibilities with respect to binding to the nsp8-nsp9-nsp10-nsp12 polyprotein. We concluded that only coordination of nsp16 following existing X-ray crystal structures (6W4H)(60) had the potential to satisfy the constraints of linking all the subunits. We then coordinated nsp14 and a second subunit of nsp10 to nsp12, following our previous model for the full hexameric replication complex (see Figure S16c).(31)

These two large complexes (nsp4-nsp5-nsp6-nsp7-nsp8/nsp4-nsp5-nsp6-nsp7/nsp13 and nsp8-nsp9-nsp10-nsp12/nsp10/nsp14/nsp16) were reasonably consistent with each other. We found they could be easily linked and optimized, connecting nsp7 to nsp8, nsp12 to nsp13 and nsp13 to nsp14. This meant only nsp9 was needed to link nsp8 and nsp10 of pp1a’ and nsp15 was needed to link nsp14 and nsp16 of pp1ab’ (see Figure S16d). Protein-protein docking with Piper(62) was used to explore initial placement of the final proteins. The most promising structures were then optimized with constraints to minimize the linkage distances before finally linking the completed polyproteins and doing a refinement of the full protein heterodimer. In the case of the nsp14-nsp15 and nsp15-nsp16 linkage, significant freedom was given to nsp9-nsp10/nsp16 to adopt a conformation that would facilitate the final linkage. The final model is shown in Figure 4c. While the conformational space of the two polyproteins is enormous and could not be exhaustively studied, in the end, we established that multiple known protein-protein interactions could be preserved in the polyprotein heterodimer. These include: pp1a’-nsp8/pp1ab’-nsp7; pp1a’-nsp8/pp1ab’-nsp13; pp1a’-nsp10/pp1ab’-nsp14; pp1ab’-nsp8/pp1ab’-nsp12; and pp1ab’-nsp10/pp1ab’-nsp16. This heterodimer was then incorporated into the dodecameric model where it went through an additional round of refinement to remove any clashes between the various components (Figure 4d).

### SARS-CoV-2 polyprotein cleavage

The close-packed arrangement of nsp5 in the polyprotein dodecameric ring establishes a favorable environment for the protease to make the initial nsp4-nsp5 cleavage. A key observation we made was that the nsp4-nsp5 linker (residues 488-510) was long enough that it could conceivably connect nsp4 to either of two nsp5 subunits in the ring (at a distance of 35 Å or 52 Å). As originally constructed from the AlphaFold hexamer, the connection was to the nearer nsp5-nsp6 subunit, comparable to how the proteins are arranged in the monomeric polyprotein. However, connecting nsp4 instead to the neighboring nsp5-nsp6 subunit allowed the nsp4-nsp5 cleavage site to pass over the proximal nsp5 protease active site. A positional shift of the nsp5 proteins would still be required for cleavage to occur, and this was investigated for both subunits. We found that simply by shifting alternating nsp5 subunits while retaining the positions of everything else, we could identify a conformation that was capable of nsp4-nsp5 cleavage, consistent with available crystal structures of bound substrate.(63) As depicted in Figures 5a and 5b, the shifted nsp5 subunit functions as the substrate, while the nsp5 subunit which retained its original position functions as the protease. Given the constraints imposed by the full (pp1a’/pp1ab’)_6_ model, a repositioning of nsp5 associated with pp1ab’ appears more likely than that of pp1a’ suggesting the larger polyprotein is the initial substrate for cleavage.

**Figure 5.**
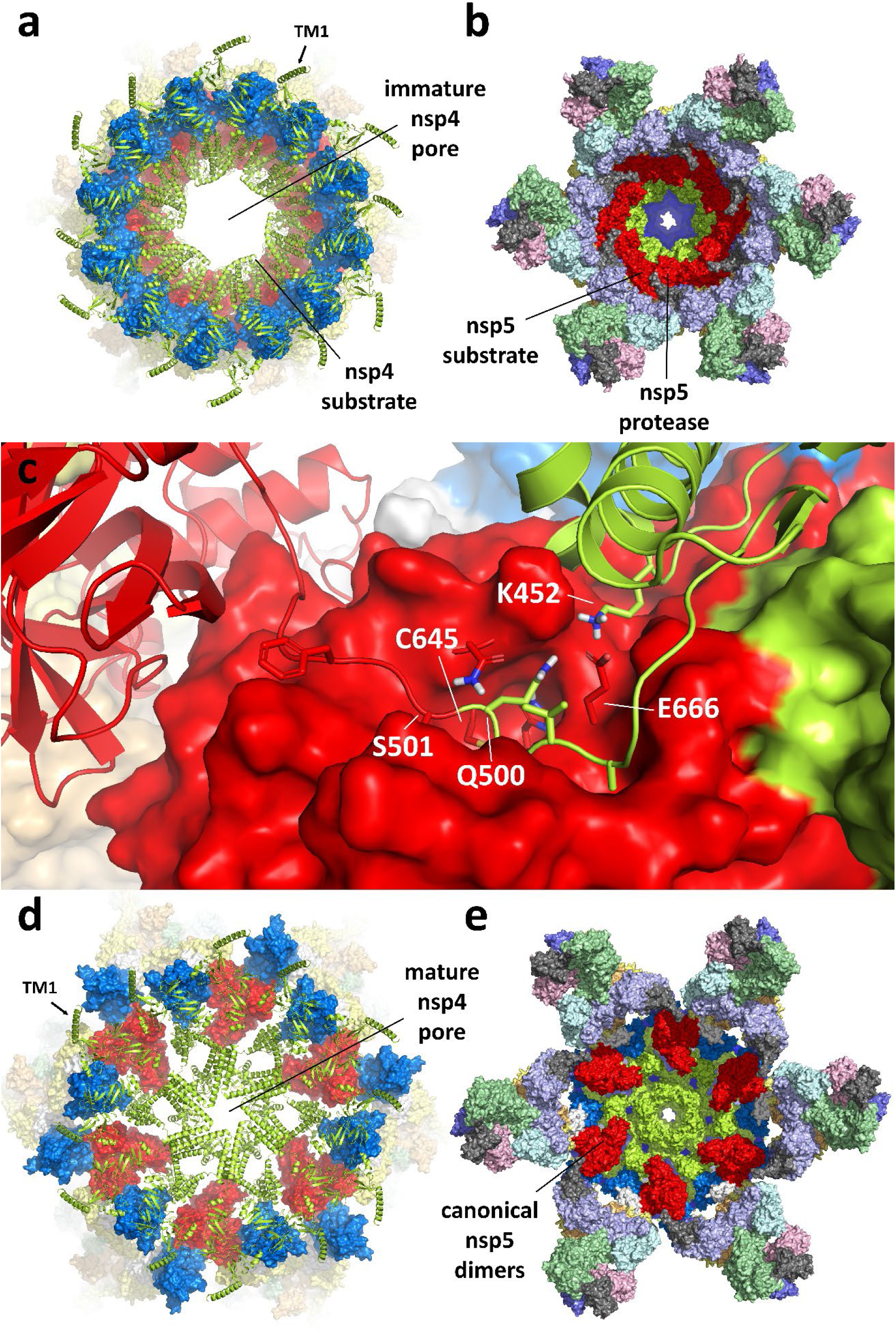
Proposed mechanism for nsp5 protease activation and pore maturation. a) View from above and b) view from below the immature pore in its pre-cleavage state. Six alternating nsp5 subunits within the dodecamer formed from pp1a’ and pp1ab’ assume a conformation which allows *trans*-cleavage of the nsp4-nsp5 linker by the remaining nsp5 subunits. c) Detail of the nsp5 protease active site shows cleavage occurs between Q500 on nsp4 and S501 on nsp5 (numbering consistent with the uncleaved polyprotein pp1ab’). The interaction between nsp4 K452 and nsp5 E666 mimics the activating interaction between the nsp5 N-terminus and the same glutamic acid residue in the canonical nsp5 dimer. d) View from above and e) view from below the post-cleavage mature pore. The reduced structural constraints from the nsp4-nsp5 cleavage of ppa1b’ allows the nsp4 subunits to adopt the conformation seen in the structure by Huang, et al.(14) Despite pp1a’ remaining uncleaved at this point, canonical nsp5 dimers can form in which one of the two protease active sites should be considered active and well positioned to cleave additional linkages.

As previously described, the protease activity of nsp5 is known to be significantly enhanced when the protein is in its canonical dimer form, in large part due to stabilization of the S1 pocket by the N-terminal amine of the paired subunit.(52) A striking observation from these AlphaFold derived models was that the C-terminal domain of nsp4 forms an interaction with nsp5 which appears to mimic this aspect of nsp5 dimerization (Figure 5c). Specifically, a conserved nsp4 basic residue (K452) forms a salt bridge with the conserved nsp5 glutamic acid (E666), replicating the interaction seen in the dimer between the same glutamic acid and the N-terminus of the paired subunit. The similar interaction seen here suggests formation of the pore prior to polyprotein cleavage both activates the nsp5 protease and positions the nsp4-nsp5 substrate to carry out the first cleavage event.

This proposed arrangement of alternating positions of nsp5 in the dodecameric ring would lead to the cleavage of half of the nsp4-nsp5 linkages. Once this occurs, the reduced constraints allow the nsp3/nsp4 pore to adopt a conformation consistent with that of the cryo-ET structure. That is, the six cleaved nsp4 proteins rotate inward to form the central hexameric pore. With this reconfiguration, the six remaining uncleaved nsp4 subunits can rotate outward to adopt the conformations seen in the cryo-ET structure. At the same time, the six partially cleaved nsp5 subunits can now form canonical nsp5 dimers with the six uncleaved nsp5 subunits, as shown in Figures 5d and 5e. In this case, the protease site in the uncleaved nsp5 subunit would be considered activated by the N-terminal amine from the cleaved subunit, as previously described. From this configuration, cleavage of additional linkages could take several pathways, with nsp5-nsp6, nsp6-nsp7, and nsp8-nsp9 linkages all within immediate range (Figure S17).

## Discussion

The events that lead to nsp5 protease cleavage of the coronavirus ORF1 polyproteins pp1a and pp1ab are at present ill-defined. Furthermore, there is currently no clear hypothesis for how the resulting cleavage products, which constitute the replication machinery, are recruited to the closed DMV replication organelle. Cleavage of nsp1, nsp2 and nsp3 from the polyprotein occurs very quickly, facilitated by the nsp3 PL proteases(17). Cleavage of the remainder of the polyprotein by nsp5 appears to be a slower process, possibly requiring dimer formation or some other regulatory event to initiate(64). As part of an investigation into the structure of the SARS-CoV-2 DMV nsp3/nsp4 pore, we were struck by a particular AlphaFold generated model of a ring formed by a multimer of uncleaved nsp4-nsp5-nsp6 polyproteins. We considered the possibility that such a structure may represent a precursor to formation of the DMV pore in an infected cell. From this starting point, we generated a series of structural models outlining a plausible path for DMV and pore formation stemming from the interaction of twelve nsp3 subunits with six pp1a’ and six pp1ab’ uncleaved polyproteins. We argue that it is the formation of an immature pore from these components that initiates the protease activity of nsp5 and creates a situation where the cleaved products would be positioned within the DMV prior to its closure.

We made several key observations from this modeling effort. First, analysis of the nsp4-nsp5-nsp6-nsp7 monomer made clear that *cis*-cleavage of the nsp4-nsp5 and nsp5-nsp6 linkages is not possible, implying cleavage is facilitated by some interaction between polyproteins. Based on a great deal of biochemical and crystallographic work on cleaved constructs,(18, 19, 21, 50, 52, 54) it is well established that nsp5 is significantly more active as a dimer. But further analysis of the nsp4-nsp5-nsp6-nsp7 dimer indicated that the constraints imposed by the flanking membrane binding proteins nsp4 and nsp6 effectively render the canonical nsp5 dimer inactive within the context of the polyprotein. Such a dimer can only be considered active following nsp4-nsp5 cleavage of at least one of the two subunits. This leads to the conclusion that some alternative interaction between the polyproteins is responsible for the initial nsp4-nsp5 cleavage event.

Second, the uncleaved polyproteins pp1a’ and pp1ab’ can form a favorable heterodimer in which known protein-protein interactions are established. While pp1a’ and pp1ab’ homodimers could also be imagined, neither would have the advantages of the complementarity seen in the heterodimer. As driven by a conformational change in the lumenal domain of nsp4, the pp1a’/pp1ab’ dimer is capable of binding to either a single membrane or a double membrane. When bound to the single membrane of the ER, the nsp4 N-terminus and CTD, nsp5, nsp7-nsp10 of pp1a’ and nsp7-nsp16 of pp1ab’ are cytosolic, while only the nsp4 lumen domain is in the ER interior. When bound to the double membrane of a DMV, the nsp4 N-terminus is the only element remaining exposed to the cytosol. With the nsp4 lumen domain sitting in the membrane gap, the bulk of the polyprotein dimer then resides in the DMV interior.

These heterodimers, when in the double membrane spanning conformation, are then capable of forming a dodecameric ring comprised of alternating pp1a’ and ppa1b’ polyproteins. This ring has a diameter comparable to that seen for the nsp4 dodecamer in the recent nsp3/nsp4 pore structure reported by Huang, et al.(14) and can associate with the nsp3 dodecamer in the same manner. Thus, we propose that this (nsp3)_12_(pp1a’)_6_(pp1ab’)_6_ complex represents an immature form of the pore.

The polyprotein dodecameric ring leads to close-packing of nsp5 and the nsp4 CTD, providing the necessary scaffolding to facilitate *trans*-cleavage of the nsp4-nsp5 linkage. We identified a conformation within this framework in which the 23-residue nsp4-nsp5 linker of alternating subunits engages the proximal nsp5 protease active site. Critically, we also noticed that a conserved basic residue from the nsp4 C-terminal domain (K452) forms a salt bridge with the nsp5 glutamate residue (E666) located in the protease active site S1 pocket. This interaction mimics the situation in the nsp5 dimer, in which the same glutamate residue interacts with the N-terminal amine of the other subunit. It is thought to be critical to stabilization of the protease active site, explaining the significant enhancement of protease activity associated with dimerization(54). Thus, we conclude that the formation of this immature pore activates the nsp5 protease and sets it up for the initial nsp4-nsp5 cleavage.

Following this initial nsp4-nsp5 cleavage event, the six cleaved nsp4 proteins and the six still uncleaved nsp4 proteins are free to adopt the conformations seen in the mature pore structure. The reduced structural constraints also allow formation of canonical nsp5 dimers in which one of the two protease sites would be activated. With these nsp5 dimers having proximity to additional cleavage sites within the polyprotein, the remainder of the polyprotein can be processed, leading to the final fully cleaved (nsp3)_12_(nsp4)_12_ pore.

With this model, we found that pore formation prior to nsp5 polyprotein processing is a reasonable possibility and may in fact be the regulatory step needed to ensure proper localization of the cleaved proteins that make up the replication machinery. A notable observation of the full model presented here is that the stoichiometry is consistent with our previous model for the hexameric replication complex(31). That model included 60+ subunits centered around nsp15. Specifically, the model includes six subunits of nsp7, nsp12, nsp13, nsp14, nsp15, and nsp16, and twelve subunits of nsp8 and nsp10, with up to six transiently associated nsp9 subunits. This complex is most efficiently derived from six pp1a and six pp1ab polyproteins, meaning the precursor presented here simultaneously produces one pore structure and generates one full replication complex. When viewed together, the two models imply a mechanism not only for DMV and pore formation, but for recruitment of the replicase to the DMV.

As depicted in Figure 6, the following picture emerges from these models. Upon cleavage by the PL proteases, nsp3 then associates with the nsp4 component of either pp1a’ or pp1ab’ on the ER membrane (Figure 6a). The nsp3 3Ecto and nsp4 lumenal domains are in the ER lumen, with the bulk of the proteins exposed to the cytosol. The association between nsp3 and nsp4 then induces double membrane formation, with aggregation into a tetrameric unit (nsp3)_2_(pp1a’)(pp1ab’) following (Figure 6b). The exact sequence of events is unclear, but this pathway seems likely given the immediate proximity between nsp3 and either pp1a’ or pp1ab’ following PLpro cleavage and the observation that the nsp3/nsp4 interaction drives membrane rearrangement.(55) Once the double membrane forms, the nsp3 3Ecto and the nsp4 lumen domains are positioned in the gap between the two membranes, and the remainder of nsp3 is on one side of the double membrane and nsp5, nsp7 and the rest of the polyprotein are on the other side. Further aggregation of these tetrameric building blocks leads to the immature pore and may facilitate the observed increased curvature of the double membrane (Figure 6c). Formation of additional pores will ultimately yield a sealed, spherical DMV.

**Figure 6.**
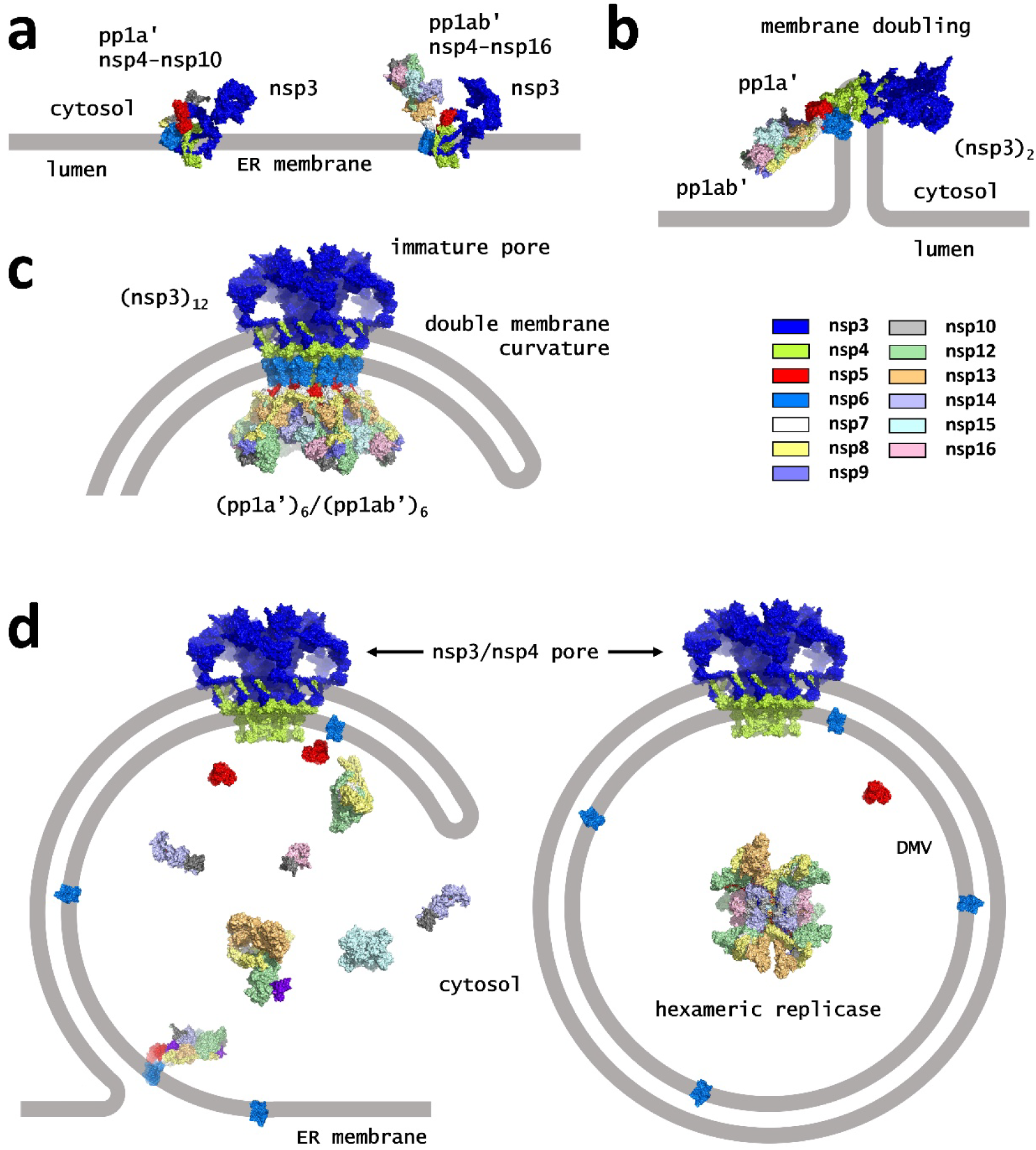
Schematic of pore formation, membrane remodeling and polyprotein cleavage. a) Following protease cleavage by PLpro, nsp3 and the polyprotein intermediates pp1a’ (nsp4-nsp10) and ppa1b’ (nsp4-nsp16) are expressed on the ER membrane. Given their initial proximity, nsp3/pp1a’ and nsp3/pp1ab’ dimers may form first. b) Interaction between nsp3 and nsp4 leads to a conformational change in the nsp4 lumen domain which drives membrane doubling, while association between pp1a’ and pp1ab’ creates the basic (nsp3)_2_(pp1a’)(pp1ab’) building block for construction of the pore. c) Further aggregation leads to formation of the dodecameric pore precursor and membrane curvature. The close packing of nsp5 in this complex activates its protease. d) Cleavage of the polyprotein prior to DMV closure releases viral proteins into the cytosol (left). Cleavage following DMV closure traps the viral proteins within the DMV, creating ideal conditions for the previously predicted hexameric replicase to form.(31)

The picture of membrane remodeling presented here raises the question of the relative timing of nsp5 polyprotein processing and DMV formation. We see no justification for a strict requirement that a DMV be fully formed for cleavage to begin. Should cleavage occur prior to DMV formation, the nsp products would be released into the cytosol. We note that in this situation, cleaved nsp6 could migrate back to the ER surface where it would be available for membrane zippering, a key step in forming the connectors for DMV clusters.(10) However, should cleavage follow closure of the DMV, the resulting products would be trapped inside (Figure 6d). This is a scenario that would likely follow formation of the final pore of the DMV, suggesting each DMV contains at minimum one hexameric replication complex. Interestingly, this process may ultimately become self-limiting, as free nsp5 dimers build up in the cytosol, cleavage of new polyproteins at late stages may occur without the requirement of new pore formation.

The proposed mechanism of regulated protease activation to ensure proper localization of the viral proteins is similar to the well-established mechanism of HIV protease maturation.(65, 66) In that case, the protease resides on the gag-pol polyprotein and requires dimerization to be enzymatically active. While the cleaved protease rapidly dimerizes in a biochemical setting, dimerization and protease activation of the polyprotein only occur within the immature viral particle. This ensures that the products (which include the reverse transcriptase and the integrase) are contained within the virion and not released into the cytosol.

The implications of this model require careful consideration. Within the protected environment of the DMV, the proposed stoichiometry, pre-organization and high protein concentration (we estimate at least 1-2 µM based on a DMV diameter of 250-350 nm(11, 12, 67)) would be ideal for formation of a long-lived full hexameric replicase complex. As previously described, this complex should ensure that replication, proofreading, capping and dsRNA unwinding are coordinated and efficient processes.(31) Confirmation of the exact contents of DMVs has been notoriously difficult, however. Multiple studies have shown that the DMVs contain RNA, specifically identifying both dsRNA and newly synthesized RNA.(3, 7, 8, 11, 67) Identification of viral protein components of the replicase within DMVs has been less conclusive, although increasing evidence supports their association with DMVs.(7, 8)

In contrast, the less controlled environment outside of DMVs may mean the full complex does not form or that complexes are transient. The minimally known replication competent complex is formed from nsp12, nsp7 and two subunits of nsp8.(68, 69) Association with nsp9(58, 70) and two subunits of nsp13(26, 57) have been documented as well, but characterization of other viral protein interactions has proven more elusive. Indeed, the proposed dual role of nsp15 suggests a full hexameric complex forms within the DMV but likely does not in the cytosol.

Within the DMV, endonuclease activity would need to be regulated to prevent degradation of the RNA. This would be the case for the hexameric replication complex, where nsp15 is proposed to function primarily as scaffolding, with its endonuclease activity highly curtailed.(31) However, in the cytosol, nsp15’s endonuclease activity serves as an effective defense against the immune response by degrading cytosolic RNA,(61, 71) suggesting the it exists as a free hexamer and is not regulated by the larger replication complex.

In the absence of full complex formation, some functions, such as proofreading and mRNA capping, might be compromised or lost entirely. Yet plenty of evidence remains that some replication is occurring outside of the DMVs. At minimum, there is a temporal argument for this, in that replication necessarily precedes DMV formation.(72) However, other lines of evidence are consistent with a two-compartment model persisting throughout infection.

Notably, the groundbreaking work of Sawicki and Sawicki(73) established that coronavirus RNA replication is dependent on continuous ORF1 translation and proteolytic processing. Further work on several temperature sensitive (TS) mutations showed that synthesis of negative-sense RNA was halted at higher temperatures but positive-sense RNA was not.(74) Donaldson, et al.(74) showed that this was a result of inhibited nsp5 protease activity at the higher temperatures. More recently, Schmidt, et al.(75) demonstrated that negative strand synthesis, was dependent on UMPylated-nsp9 priming, facilitated by the host factor SND1. This did not appear to be the case for positive strand synthesis. With no proposed mechanism for recruitment of additional proteins to an already formed DMV, the dependence of negative-sense RNA synthesis on host factors and the continuous generation of viral replicase proteins (specifically nsp9) suggest transcription of the minus strand occurs exclusively in the cytosol. In contrast, this model would be consistent with positive strand synthesis occurring in the cytosol at early stages but predominantly occurring within the protected environment of DMVs at later stages.

In summary, in asking the question whether the nsp3/nsp4 DMV pore forms prior to nsp5 cleavage of the polyprotein, we have developed a simple framework for understanding how the replication complex is recruited to DMVs in coronavirus infected cells. Our model directly links the dodecameric nsp3/nsp4 pore structure and the previously proposed 60+ subunit hexameric replication complex to the same polyprotein intermediate, elucidating how the nsp5 protease is regulated to ensure proper localization of the replicase in the DMV. From a close examination of structural constraints within monomeric and multimeric forms of the polyprotein, aided in large part by AlphaFold predictions, we conclude that formation of an immature pore from twelve nsp3 subunits, and six subunits each of the intermediate polyproteins pp1a’ and pp1ab’ initiates the protease activity of nsp5. Following cleavage of half of the nsp4-nsp5 linkages, the mature nsp3/nsp4 pore can form, and protease cleavage of the remainder of the polyproteins will release the components necessary to form the full replication complex into the DMV. While the detailed mechanism is specific to coronaviruses, we expect similar processes are taking place with other single-stranded (+) RNA viral families.

## Materials and Methods

AlphaFold2.1(76) (with a template cutoff date of 2/15/2021) was used to predict the structures of individual monomers of nsp3, nsp4, and uncleaved nsp4-nsp5-nsp6-nsp7 from SARS-CoV-2, as well as hexameric complexes of the C-terminal Y1/CoV-Y domain of nsp3, hexameric complexes of the TMD2/CTD domains of nsp4 (residues 256-500), hexameric complexes of uncleaved nsp4-nsp5-nsp6 (residues 256-1042), dimeric complexes of uncleaved nsp4-nsp5-nsp6-nsp7, and various fragments and heterodimers of nsp3 and the pp1a’ and pp1ab’ polyproteins. The SARS-CoV-2 sequences used are from the original Wuhan strain (NCBI accession number QHD43415.1(77)). Nsp3 spans residues 819-2763 of the polyprotein, nsp4 spans 2764-3263, pp1a’ (nsp4-nsp10) spans 2764-4392, and pp1ab’ (nsp4-nsp16) spans 2764-7096. The MHV sequence used in construction of the nsp3 dodecamer was from NCBI sequence AAX23975.1.(78) Additional detail on the constructs explored can be found in the Supplementary Information.

In all cases, AMBER relaxation was turned off and structures were instead refined with software from the Schrӧdinger Suite, including Prime, Macromodel and Desmond(79). In all cases, the OPLS4 force field was employed.(80) A typical workflow involved construction of an initial model either from AlphaFold directly or via protein-protein docking with Piper(81) or manual manipulation of individual subunits, followed by sidechain prediction in Prime. This was followed by a stepwise minimization with Prime and/or Macromodel, in which residues at the protein-protein interface were optimized first, building up to a minimization of the full structure. In cases where large movements from the starting point occurred, a second round of optimization would be carried out starting from a structure in which the original protein subunits were re-aligned to their new positions. Stability was tested at several stages of model development with molecular dynamics (MD) using Desmond, typically run for 100ns at constant temperature and pressure (NPT) with suitable constraints. SPC water was used in these simulations, as well as a POPC membrane, if appropriate. Macromodel and Desmond were also used to predict conformations of some long loops. Construction of the pp1a’/pp1ab’ dimer involved preservation of known interactions as much as possible, using AlphaFold and Piper protein-protein docking to explore possible arrangements of various subunits. Model quality was assessed with Schrödinger’s Protein Reports and by MolProbity.(82)

Models were guided by the cryo-ET maps deposited in the Electron Microscopy Data Bank with accession codes EMD-11514 (MHV(12)), EMD-15963 (SARS-CoV-2(13)), and EMD-39107 and EMD-39109 (SARS-CoV-2(14)). Ultimately, the models were merged with the structures of Huang, et al. (PDB 8YAX and 8YB5)(14) and further refined using the approaches described above. While the final pore models have approximate C6 rotational symmetry, strict symmetry constraints were not imposed.

## Data Availibility

All data needed to evaluate the conclusions of this work are present in the paper and/or the Supplementary Materials. Structure models in CIF format are available upon request.

## Notes

### Competing Interest Statement

The authors have declared no competing interest.

### Summary of Updates

The revised manuscript includes a significant amount of additional detail, including how the models were constructed and validated. Notably, we show how formation of the canonical nsp5 dimer within the context of the uncleaved polyprotein cannot be responsible for the initial cleavage event. We also include a control study showing how a similar modeling approach accurately reproduces the cryo-EM structure of the chikungunya virus nsP1 dodecamer.

