## Supplementary Information and Figures for "A structural roadmap for the formation of the coronavirus nsp3/nsp4 double membrane vesicle pore and its implications for polyprotein processing and replication/transcription"

**Supplementary Results**

*SARS-CoV-2 nsp3 dodecamer detail*

As described in the primary text, portions of the nsp3 dodecamer component of the cryo-ET pore structure from Huang, et al.(1) were remodeled. The resolution of the prongs was considerably weaker (> 9 Å) than that of the crown but was fit by Huang, et al. with the Mac2, Mac3, DPUP and NAB domains nonetheless, relying on an Alphafold prediction of the Mac2/Mac3/NAB complex and docking of that model to the density. Not fit to any density were the Ubl1, HVR and Mac1 domains, despite previous work by Zimmermann, et al.(2), demonstrating that these domains were major contributors to the bulk and cohesion of the prong. Also not fit to any density was the βSM domain, which is predicted by AlphaFold to be connected to TM1 via a pair of amphipathic helices. This would position the domain at the base of the crown, an assumption also suggested by Huang, et al. A particular challenge then for the published fit to the prongs is that the NAB domain must connect via a 44-residue linker to the βSM domain. While technically this may be possible, the linker would need to be nearly straight chain to achieve this (we estimated a distance of at least 130 Å between the two domains, whereas a perfectly straight chain 44 amino acid peptide would be 160 Å).

In examination of the cryo-ET maps, we identified previously unfit density stemming from TM1 which was consistent with amphipathic helices running along the cytosolic face of the outer membrane. As shown in Figure S4a, we found we could fit the predicted amphipathic helices connecting βSM and TM1 in this density for six subunits. This fit placed the βSM domain at the base of the crown, albeit without clear density to confirm the exact orientation. In this case, the lower resolution map for the MHV pore(3) provided additional constraints on the placement of the conserved βSM domain.

A factor in refitting the prongs was consideration of sequence conservation across betacoronaviruses, reasoning that conserved domains would serve to anchor the prongs to the crown and the more unique domains would be in distal positions. The MHV map was helpful in this regard. Overall, MHV has a different shape to the prongs, reflecting differences among the N-terminal domains. While the Ubl1, DPUP and NAB domains are preserved, and the MHV Mac domain corresponds to the SARS-CoV-2 Mac1 domain, MHV has an additional papain-like protease (PLP1) and does not have either Mac2 or Mac3 domains. The N-terminal domains are also arranged differently, Ubl1-HVR-PLP1-Mac-DPUP for MHV and Ubl1-HVR-Mac1-Mac2-Mac3-DPUP for SARS-CoV-2. Wolff, et al.(3) demonstrated that the MHV N-terminal Ubl1 domain sits at the tip of the prong, implying that the HVR and PLP1 domains occupy the bulk of the prong, connecting to the rim of the crown via the Mac and DPUP domains. Given that the Mac (Mac1) domain is important enzymatically, containing the ADP-ribose phosphatase (ADRP) active site(4), we reasoned that this domain should reside in a similar location for both MHV and SARS-CoV-2. We also reasoned that it would necessarily be in proximity to the DPUP domain based on the direct linkage between these two domains in MHV. Further functional constraints used in placement of the N-terminal domains included the Mac2 domain being in a position suitable to interact with the host translation co-factor Paip1(5) and the NAB domain exposing its conserved RNA-binding residues.

Ultimately, we fit the NAB domain in the rim of the crown between neighboring PLpro domains, while the DPUP and Mac1 domains were positioned to form the base of the prong. Ubl1, Mac2 and Mac3 were then used to fill out the remainder of the prong. As shown in Figure S4b, these domains offer a superior fit to the prong density. The density is more ambiguous for placement of the NAB domain, although the lower resolution maps would suggest that space is occupied. Importantly, the NAB domain is nearly 50 Å closer to the βSM domain (91 Å vs. 138 Å as measured from residue 1194 to 1242), allowing them to be reasonably connected, as we have done as a proof of concept. Furthermore, the basic residues that constitute the RNA binding motif of NAB are well exposed (Figure S4c), positioned along the inner rim of the crown such that they may direct RNA ejected from the pore toward the prongs. In Figure S4d, we show that Paip1 is capable of binding to Mac2 on the top of the prong, while the ADPR active site is well exposed at the bottom of the prong.

Consistent with this model, we constructed a model for the MHV nsp3 dodecameric crown. In Figure S5, we compare our model of the SARS-CoV-2 prong in the Zimmermann, et al. map (Figure S5a) vs. our model of the MHV prong in the Wolff, et al. map (Figure S4b). In Figure S5c, we show a detailed overlay of the two models, focusing on the common domains that make up the rim of the crown and the base of the prongs. Note that the published Zimmerman map appears to be inverted. The handedness of cryo-ET reconstructions may be flipped at multiple stages of data collection and reconstruction. Without achieving a resolution high enough to visualize the orientation of helices or utilizing protocols with specialized fiducial markers(6), the true handedness of the reconstruction may be ambiguous. As the Huang, et al. structure resolved the ambiguity, we generated a mirror image of the Zimmerman map, which proved to be consistent both with the Huang map and the Wolff map.

While the models presented here went through multiple rounds of conformational sampling and energy minimization, we stress that the SARS-CoV-2 model is built on a variety of constraints and does not represent a strict re-refinement of the cryo-ET structure.

### Supplementary Methods

*Alphafold constructs*

SARS-CoV-2 nsp3 (QHD43415.1):(7)

APTKVTFGDDTVIEVQGYKSVNITFELDERIDKVLNEKCSAYTVELGTEVNEFACVVADAVIKTLQPVSELLTPL
GIDLDEWSMATYYLFDESGEFKLASHMYCSFYPPDEDEEEGDCEEEEFEPSTQYEYGTEDDYQGKPLEFGATS
AALQPEEEQEEDWLDDDSQQTVGQQDGSEDNQTTTIQTIVEVQPQLEMELTPVVQTIEVNSFSGYLKLTDN
VYIKNADIVEEAKKVKPTVVVNAANVYLKHGGGVAGALNKATNNAMQVESDDYIATNGPLKVGGSCVLSG
HNLAKHCLHVVGPNVNKGEDIQLLKSAYENFNQHEVLLAPLLSAGIFGADPIHSLRVCVDTVRTNVYLAVFDK
NLYDKLVSSFLEMKSEKQVEQKIAEIPKEEVKPFITESKPSVEQRKQDDKKIKACVEEVTTTLEETKFLTENLLLYI
DINGNLHPDSATLVSDIDITFLKKDAPYIVGDVVQEGVLTAVVIPTKKAGGTTEMLAKALRKVPTDNYITTYPG
QGLNGYTVEEAKTVLKKCKSAFYILPSIISNEKQEILGTVSWNLREMLAHAEETRKLMPVCVETKAIVSTIQRKY
KGIKIQEGVVDYGARFYFYTSKTTVASLINTLNDLNETLVTMPLGYVTHGLNLEEAARYMRSLKVPATVSVSSP
DAVTAYNGYLTSSSKTPEEHFIETISLAGSYKDWSYSGQSTQLGIEFLKRGDKSVYYTSNPTTFHLDGEVITFDN
LKTLLSLREVRTIKVFTTVDNINLHTQVVDMSMTYGQQFGPTYLDGADVTKIKPHNSHEGKTFYVLPNDDTLR
VEAFEYYHTTDPSFLGRYMSALNHTKKWKYPQVNGLTSIKWADNNCYLATALLTLQQIELKFNPPALQDAYY
RARAGEAANFCALILAYCNKTVGELGDVRETMSYLFQHANLDSCKRVLNVVCKTCGQQQTTLKGVEAVMY
MGTLSYEQFKKGVQIPCTCGKQATKYLVQQESPFVMMSAPPAQYELKHGTFTCASEYTGNYQCGHYKHITS
KETLYCIDGALLTKSSEYKGPITDVFYKENSYTTTIKPVTYKLDGVVCTEIDPKLDNYYKKDNSYFTEQPIDLVPN
QPYPNASFDNFKFVCDNIKFADDLNQLTGYKKPASRELKVTFFPDLNGDVVAIDYKHYTPSFKKGAKLLHKPIV
WHVNNATNKATYKPNTWCIRCLWSTKPVETSNSFDVLKSEDAQGMDNLACEDLKPVSEEVVENPTIQKDV
LECNVKTTEVVGDIILKPANNSLKITEEVGHTDLMAAYVDNSSLTIKKPNELSRVLGLKTLATHGLAAVNSVPW
DTIANYAKPFLNKVVSTTTNIVTRCLNRVCTNYMPYFFTLLLQLCTFTRSTNSRIKASMPTTIAKNTVKSVGKFC
LEASFNYLKSPNFSKLINIIIWFLLLSVCLGSLIYSTAALGVLMSNLGMPSYCTGYREGYLNSTNVTIATYCTGSIP
CSVCLSGLDSLDTYPSLETIQITISSFKWDLTAFGLVAEWFLAYILFTRFFYVLGLAAIMQLFFSYFAVHFISNSWL
MWLIINLVQMAPISAMVRMYIFFASFYYVWKSYVHVVDGCNSSTCMMCYKRNRATRVECTTIVNGVRRSF
YVYANGGKGFCKLHNWNCVNCDTFCAGSTFISDEVARDLSLQFKRPINPTDQSSYIVDSVTVKNGSIHLYFDK
AGQKTYERHSLSHFVNLDNLRANNTKGSLPINVIVFDGKSKCEESSAKSASVYYSQLMCQPILLLDQALVSDV

GDSAEVAVKMFDAYVNTFSSTFNVPMEKLKTLVATAEAELAKNVSLDNVLSTFISAARQGFVDSDVETKDVV
ECLKLSHQSDIEVTGDSCNNYMLTYNKVENMTPRDLGACIDCSARHINAQVAKSHNIALIWNVKDFMSLSE
QLRKQIRSAAKKNNLPFKLTCATTRQVVNVVTTKIALKGG

Constructs include: full-length, Ubl1-HVR-Mac1 (1-387), Mac1-Mac2 (212-539), Mac2-Mac3 (416-674), Mac3-DPUP (547-744), DPUP-Ubl2/PLpro (678-1093), Ubl2/PLpro-NAB (747-1196), NAB-βSM (1109-1365), βSM-TM1/3Ecto/TMD2-Y1 (1241-1761), Y1/CoV-Y (1596-1945), Mac1-Mac3 (212-674), Mac2 (413-549), Mac2-Mac3 (413-674), hexameric Y1/CoV-Y (1587-1945)

SARS-CoV-2 nsp4 full-length (QHD43415.1):

KIVNNWLKQLIKVTLVFLFVAAIFYLITPVHVMSKHTDFSSEIIGYKAIDGGVTRDIASTDTCFANKHADFDTWF
SQRGGSYTNDKACPLIAAVITREVGFVVPGLPGTILRTTNGDFLHFLPRVFSAVGNICYTPSKLIEYTDFATSAC
VLAAECTIFKDASGKPVPYCYDTNVLEGSVAYESLRPDTRYVLMDGSIIQFPNTYLEGSVRVVTTFDSEYCRHG
TCERSEAGVCVSTSGRWVLNNDYYRSLPGVFCGVDAVNLLTNMFTPLIQPIGALDISASIVAGGIVAIVVTCLA
YYFMRFRRAFGEYSHVVAFNTLLFLMSFTVLCLTPVYSFLPGVYSVIYLYLTFYLTNDVSFLAHIQWMVMFTPL
VPFWITIAYIICISTKHFYWFFSNYLKRRVVFNGVSFSTFEEAALCTFLLNKEMYLKLRSDVLLPLTQYNRYLALY
NKYKYFSGAMDTTSYREAACCHLAKALNDFSNSGSDVLYQPPQTSITSAVLQ

Constructs include: full-length, TMD2-CTD (256-500)

SARS-CoV-2 nsp4-nsp7 polyprotein (QHD43415.1):

KIVNNWLKQLIKVTLVFLFVAAIFYLITPVHVMSKHTDFSSEIIGYKAIDGGVTRDIASTDTCFANKHADFDTWF
SQRGGSYTNDKACPLIAAVITREVGFVVPGLPGTILRTTNGDFLHFLPRVFSAVGNICYTPSKLIEYTDFATSAC
VLAAECTIFKDASGKPVPYCYDTNVLEGSVAYESLRPDTRYVLMDGSIIQFPNTYLEGSVRVVTTFDSEYCRHG
TCERSEAGVCVSTSGRWVLNNDYYRSLPGVFCGVDAVNLLTNMFTPLIQPIGALDISASIVAGGIVAIVVTCLA
YYFMRFRRAFGEYSHVVAFNTLLFLMSFTVLCLTPVYSFLPGVYSVIYLYLTFYLTNDVSFLAHIQWMVMFTPL
VPFWITIAYIICISTKHFYWFFSNYLKRRVVFNGVSFSTFEEAALCTFLLNKEMYLKLRSDVLLPLTQYNRYLALY
NKYKYFSGAMDTTSYREAACCHLAKALNDFSNSGSDVLYQPPQTSITSAVLQSGFRKMAFPSGKVEGCMVQ
VTCGTTTLNGLWLDDVVYCPRHVICTSEDMLNPNYEDLLIRKSNHNFLVQAGNVQLRVIGHSMQNCVLKLK
VDTANPKTPKYKFVRIQPGQTFSVLACYNGSPSGVYQCAMRPNFTIKGSFLNGSCGSVGFNIDYDCVSFCYM
HHMELPTGVHAGTDLEGNFYGPFVDRQTAQAAGTDTTITVNVLAWLYAAVINGDRWFLNRFTTTLNDFNL
VAMKYNYEPLTQDHVDILGPLSAQTGIAVLDMCASLKELLQNGMNGRTILGSALLEDEFTPFDVVRQCSGVT
FQSAVKRTIKGTHHWLLLTILTSLLVLVQSTQWSLFFFLYENAFLPFAMGIIAMSAFAMMFVKHKHAFLCLFLL
PSLATVAYFNMVYMPASWVMRIMTWLDMVDTSLSGFKLKDCVMYASAVVLLILMTARTVYDDGARRVW
TLMNVLTLVYKVYYGNALDQAISMWALIISVTSNYSGVVTTVMFLARGIVFMCVEYCPIFFITGNTLQCIMLV
YCFLGYFCTCYFGLFCLLNRYFRLTLGVYDYLVSTQEFRYMNSQGLLPPKNSIDAFKLNIKLLGVGGKPCIKVAT
VQSKMSDVKCTSVVLLSVLQQLRVESSSKLWAQCVQLHNDILLAKDTTEAFEKMVSLLSVLLSMQGAVDINK
LCEEMLDNRATLQ

Constructs include: full-length, full-length dimer, hexameric truncated nsp4-nsp6 (256-1042)

SARS-CoV-2 nsp4-nsp10 polyprotein (QHD43415.1):

KIVNNWLKQLIKVTLVFLFVAAIFYLITPVHVMSKHTDFSSEIIGYKAIDGGVTRDIASTDTCFANKHADFDTWF
SQRGGSYTNDKACPLIAAVITREVGFVVPGLPGTILRTTNGDFLHFLPRVFSAVGNICYTPSKLIEYTDFATSAC
VLAAECTIFKDASGKPVPYCYDTNVLEGSVAYESLRPDTRYVLMDGSIIQFPNTYLEGSVRVVTTFDSEYCRHG
TCERSEAGVCVSTSGRWVLNNDYYRSLPGVFCGVDAVNLLTNMFTPLIQPIGALDISASIVAGGIVAIVVTCLA
YYFMRFRRAFGEYSHVVAFNTLLFLMSFTVLCLTPVYSFLPGVYSVIYLYLTFYLTNDVSFLAHIQWMVMFTPL
VPFWITIAYIICISTKHFYWFFSNYLKRRVVFNGVSFSTFEEAALCTFLLNKEMYLKLRSDVLLPLTQYNRYLALY
NKYKYFSGAMDTTSYREAACCHLAKALNDFSNSGSDVLYQPPQTSITSAVLQSGFRKMAFPSGKVEGCMVQ
VTCGTTTLNGLWLDDVVYCPRHVICTSEDMLNPNYEDLLIRKSNHNFLVQAGNVQLRVIGHSMQNCVLKLK
VDTANPKTPKYKFVRIQPGQTFSVLACYNGSPSGVYQCAMRPNFTIKGSFLNGSCGSVGFNIDYDCVSFCYM
HHMELPTGVHAGTDLEGNFYGPFVDRQTAQAAGTDTTITVNVLAWLYAAVINGDRWFLNRFTTTLNDFNL
VAMKYNYEPLTQDHVDILGPLSAQTGIAVLDMCASLKELLQNGMNGRTILGSALLEDEFTPFDVVRQCSGVT
FQSAVKRTIKGTHHWLLLTILTSLLVLVQSTQWSLFFFLYENAFLPFAMGIIAMSAFAMMFVKHKHAFLCLFLL
PSLATVAYFNMVYMPASWVMRIMTWLDMVDTSLSGFKLKDCVMYASAVVLLILMTARTVYDDGARRVW
TLMNVLTLVYKVYYGNALDQAISMWALIISVTSNYSGVVTTVMFLARGIVFMCVEYCPIFFITGNTLQCIMLV
YCFLGYFCTCYFGLFCLLNRYFRLTLGVYDYLVSTQEFRYMNSQGLLPPKNSIDAFKLNIKLLGVGGKPCIKVAT
VQSKMSDVKCTSVVLLSVLQQLRVESSSKLWAQCVQLHNDILLAKDTTEAFEKMVSLLSVLLSMQGAVDINK
LCEEMLDNRATLQAIASEFSSLPSYAAFATAQEAYEQAVANGDSEVVLKKLKKSLNVAKSEFDRDAAMQRKL
EKMADQAMTQMYKQARSEDKRAKVTSAMQTMLFTMLRKLDNDALNNIINNARDGCVPLNIIPLTTAAKL
MVVIPDYNTYKNTCDGTTFTYASALWEIQQVVDADSKIVQLSEISMDNSPNLAWPLIVTALRANSAVKLQNN
ELSPVALRQMSCAAGTTQTACTDDNALAYYNTTKGGRFVLALLSDLQDLKWARFPKSDGTGTIYTELEPPCR
FVTDTPKGPKVKYLYFIKGLNNLNRGMVLGSLAATVRLQAGNATEVPANSTVLSFCAFAVDAAKAYKDYLAS
GGQPITNCVKMLCTHTGTGQAITVTPEANMDQESFGGASCCLYCRCHIDHPNPKGFCDLKGKYVQIPTTCA
NDPVGFTLKNTVCTVCGMWKGYGCSCDQLREPMLQ

Constructs include: full-length, nsp7-nsp8 (1097-1377), nsp8-nsp9 (1180-1490), nsp9-nsp10 (1378-1629)

SARS-CoV-2 nsp4-nsp16 polyprotein (QHD43415.1):

KIVNNWLKQLIKVTLVFLFVAAIFYLITPVHVMSKHTDFSSEIIGYKAIDGGVTRDIASTDTCFANKHADFDTWF
SQRGGSYTNDKACPLIAAVITREVGFVVPGLPGTILRTTNGDFLHFLPRVFSAVGNICYTPSKLIEYTDFATSAC
VLAAECTIFKDASGKPVPYCYDTNVLEGSVAYESLRPDTRYVLMDGSIIQFPNTYLEGSVRVVTTFDSEYCRHG
TCERSEAGVCVSTSGRWVLNNDYYRSLPGVFCGVDAVNLLTNMFTPLIQPIGALDISASIVAGGIVAIVVTCLA
YYFMRFRRAFGEYSHVVAFNTLLFLMSFTVLCLTPVYSFLPGVYSVIYLYLTFYLTNDVSFLAHIQWMVMFTPL
VPFWITIAYIICISTKHFYWFFSNYLKRRVVFNGVSFSTFEEAALCTFLLNKEMYLKLRSDVLLPLTQYNRYLALY
NKYKYFSGAMDTTSYREAACCHLAKALNDFSNSGSDVLYQPPQTSITSAVLQSGFRKMAFPSGKVEGCMVQ
VTCGTTTLNGLWLDDVVYCPRHVICTSEDMLNPNYEDLLIRKSNHNFLVQAGNVQLRVIGHSMQNCVLKLK
VDTANPKTPKYKFVRIQPGQTFSVLACYNGSPSGVYQCAMRPNFTIKGSFLNGSCGSVGFNIDYDCVSFCYM
HHMELPTGVHAGTDLEGNFYGPFVDRQTAQAAGTDTTITVNVLAWLYAAVINGDRWFLNRFTTTLNDFNL
VAMKYNYEPLTQDHVDILGPLSAQTGIAVLDMCASLKELLQNGMNGRTILGSALLEDEFTPFDVVRQCSGVT
FQSAVKRTIKGTHHWLLLTILTSLLVLVQSTQWSLFFFLYENAFLPFAMGIIAMSAFAMMFVKHKHAFLCLFLL
PSLATVAYFNMVYMPASWVMRIMTWLDMVDTSLSGFKLKDCVMYASAVVLLILMTARTVYDDGARRVW

TLMNVLTLVYKVYYGNALDQAISMWALIISVTSNYSGVVTTVMFLARGIVFMCVEYCPIFFITGNTLQCIMLVYCFLGYFCTCYFGLFCLLNRYFRLTLGVYDYLVSTQEFRYMNSQGLLPPKNSIDAFKLNIKLLGVGGKPCIKVATVQSKMSDVKCTSVVLLSVLQQLRVESSSKLWAQCVQLHNDILLAKDTTEAFEKMVSLLSVLLSMQGAVDINKLCEEMLDNRATLQAIASEFSSLPSYAAFATAQEAYEQAVANGDSEVVLKKLKKSLNVAKSEFDRDAAMQRKLEKMADQAMTQMYKQARSEDKRAKVTSAMQTMLFTMLRKLDNDALNNIINNARDGCVPLNIIPLTTAAKLMVVIPDYNTYKNTCDGTTFTYASALWEIQQVVDADSKIVQLSEISMDNSPNLAWPLIVTALRANSAVKLQNNELSPVALRQMSCAAGTTQTACTDDNALAYYNTTKGGRFVLALLSDLQDLKWARFPKSDGTGTIYTELEPPCRFVTDTPKGPKVKYLYFIKGLNNLNRGMVLGSLAATVRLQAGNATEVPANSTVLSFCAFAVDAAKAYKDYLASGGQPITNCVKMLCTHTGTGQAITVTPEANMDQESFGGASCCLYCRCHIDHPNPKGFCDLKGKYVQIPTTCANDPVGFTLKNTVCTVCGMWKGYGCSCDQLREPMLQSADAQSFLNRVCGVSAARLTPCGTGTSTDVVYRAFDIYNDKVAGFAKFLKTNCCRFQEKDEDDNLIDSYFVVKRHTFSNYQHEETIYNLLKDCPAVAKHDFFKFRIDGDMVPHISRQRLTKYTMADLVYALRHFDEGNCDTLKEILVTYNCCDDDYFNKKDWYDFVENPDILRVYANLGERVRQALLKTVQFCDAMRNAGIVGVLTLDNQDLNGNWYDFGDFIQTTPGSGVPVVDSYYSLLMPILTLTRALTAESHVDTDLTKPYIKWDLLKYDFTEERLKLFDRYFKYWDQTYHPNCVNCLDDRCILHCANFNVLFSTVFPPTSFGPLVRKIFVDGVPFVVSTGYHFRELGVVHNQDVNLHSSRLSFKELLVYAADPAMHAASGNLLLDKRTTCFSVAALTNNVAFQTVKPGNFNKDFYDFAVSKGFFKEGSSVELKHFFFAQDGNAAISDYDYYRYNLPTMCDIRQLLFVVEVVDKYFDCYDGGCINANQVIVNNLDKSAGFPFNKWGKARLYYDSMSYEDQDALFAYTKRNVIPTITQMNLKYAISAKNRARTVAGVSICSTMTNRQFHQKLLKSIAATRGATVVIGTSKFYGGWHNMLKTVYSDVENPHLMGWDYPKCDRAMPNMLRIMASLVLARKHTTCCSLSHRFYRLANECAQVLSEMVMCGGSLYVKPGGTSSGDATTAYANSVFNICQAVTANVNALLSTDGNKIADKYVRNLQHRLYECLYRNRDVDTDFVNEFYAYLRKHFSMMILSDDAVVCFNSTYASQGLVASIKNFKSVLYYQNNVFMSEAKCWTETDLTKGPHEFCSQHTMLVKQGDDYVYLPYPDPSRILGAGCFVDDIVKTDGTLMIERFVSLAIDAYPLTKHPNQEYADVFHLYLQYIRKLHDELTGHMLDMYSVMLTNDNTSRYWEPEFYEAMYTPHTVLQAVGACVLCNSQTSLRCGACIRRPFLCCKCCYDHVISTSHKLVLSVNPYVCNAPGCDVTDVTQLYLGGMSYYCKSHKPPISFPLCANGQVFGLYKNTCVGSDNVTDFNAIATCDWTNAGDYILANTCTERLKLFAAETLKATEETFKLSYGIATVREVLSDRELHLSWEVGKPRPPLNRNYVFTGYRVTKNSKVQIGEYTFEKGDYGDAVVYRGTTTYKLNVGDYFVLTSHTVMPLSAPTLVPQEHYVRITGLYPTLNISDEFSSNVANYQKVGMQKYSTLQGPPGTGKSHFAIGLALYYPSARIVYTACSHAAVDALCEKALKYLPIDKCSRIIPARARVECFDKFKVNSTLEQYVFCTVNALPETTADIVVFDEISMATNYDLSVVNARLRAKHYVYIGDPAQLPAPRTLLTKGTLEPEYFNSVCRLMKTIGPDMFLGTCRRCPAEIVDTVSALVYDNKLKAHKDKSAQCFKMFYKGVITHDVSSAINRPQIGVVREFLTRNPAWRKAVFISPYNSQNAVASKILGLPTQTVDSSQGSEYDYVIFTQTTETAHSCNVNRFNVAITRAKVGILCIMSDRDLYDKLQFTSLEIPRRNVATLQAENVTGLFKDCSKVITGLHPTQAPTHLSVDTKFKTEGLCVDIPGIPKDMTYRRLISMMGFKMNYQVNGYPNMFITREEAIRHVRAWIGFDVEGCHATREAVGTNLPLQLGFSTGVNLVAVPTGYVDTPNNTDFSRVSAKPPPGDQFKHLIPLMYKGLPWNVVRIKIVQMLSDTLKNLSDRVVFVLWAHGFELTSMKYFVKIGPERTCCLCDRRATCFSTASDTYACWHHSIGFDYVYNPFMIDVQQWGFTGNLQSNHDLYCQVHGNAHVASCDAIMTRCLAVHECFVKRVDWTIEYPIIGDELKINAACRKVQHMVVKAALLADKFPVLHDIGNPKAIKCVPQADVEWKFYDAQPCSDKAYKIEELFYSYATHSDKFTDGVCLFWNCNVDRYPANSIVCRFDTRVLSNLNLPGCDGGSLYVNKHAFHTPAFDKSAFVNLKQLPFFYYSDSPCESHGKQVVSDIDYVPLKSATCITRCNLGGAVCRHHANEYRLYLDAYNMMISAGFSLWVYKQFDTYNLWNTFTRLQSLENVAFNVVNKGHFDGQQGEVPVSIINNTVYTKVDGVDVELFENKTTLPVNVAFELWAKRNIKPVPEVKILNNLGVDIAANTVIWDYKRDAPAHISTIGVCSMTDIAKKPTETICAPLTVFFDGRVDGQVDLFRNARNGVLITEGSVKGLQPSVGPKQASLNGVTLIGEAVKTQFNYYKKVDGVVQQLPETYFTQSRNLQEFKPRSQMEIDFLELAMDEFIERYKLEGYAFEHIVYGDFSHSQLGGLHLLIGLAKRFKESPFELEDFIPMDSTVKNYFITDAQTGSSKCVCSVIDLLLDDFVEIIKSQDLSVVSKVVKVTIDYTEISFMLWCKDGHVETFYPKLQSSQAWQPGVAMPNLYKMQRMLLEKCDLQNYGDSATLPKGIMMNVAKYTQLCQYLNTLTLAVPYNMRVIHFGAGSDKGVAPGTAVLRQ

WLPTGTLLVDSDLNDFVSDADSTLIGDCATVHTANKWDLIISDMYDPKTKNVTKENDSKEGFFTYICGFIQQK
LALGGSVAIKITEHSWNADLYKLMGHFAWWTAFVTNVNASSSEAFLIGCNYLGKPREQIDGYVMHANYIFW
RNTNPIQLSSYSLFDMSKFPLKLRGTAVMSLKEGQINDMILSLLSKGRLIIRENNRVVISSDVLVNN

Constructs include: full-length, nsp10-nsp12 (1491-2561), nsp12-nsp13 (1630-3162), nsp13-nsp14 (2562-3689), nsp14-nsp15 (3163-4035), nsp15-nsp16 (3690-4333), nsp8-nsp12 (1180-2561), heterodimer nsp8-nsp12 (1180-2561)/nsp14 (3163-3689), heterodimer nsp8-nsp12 (1180-2561)/nsp16 (4036-4333), heterodimer nsp8-nsp12 (1180-2561)/nsp15-nsp16 (3690-4333)

Human Paip1 (6YXJ):(5)

GSHMASMTGGQQMGRGSTLSEYVQDFLNHLTEQPGSFETEIEQFAETLNGCVTTDDALQELVELIYQQATSI
PNFSYMGARLCNYLSHHLTISPQSGNFRQLLLQRCRTEYEVKDQAAKGDEVTRKRFHAFVLFLGELYLNLEIKG
TNGQVTRADILQVGLRELLNALFSNPMDDNLICAVKLLKLTGSVLEDAWKEKGKMDMEEIIQRIENVVLDAN
CSRDVKQMLLKLVELRSS

Constructs: Mac2 (413-549)/Paip1, Mac2-Mac3 (413-674)/Paip1

MHV nsp3 (AAX23975.1):(8)

KKVEFNDKPKVRKIPSTRKIKITFALDATFDSVLSKACSEFEVDKDVTLDELLDVVLDAVESTLSPCKEHDVIGTK
VCALLDRLAGDYVYLFDEGGDEVIAPRMYCSFSAPDDEDCVAADVVDADENQDDDAEDSAVLVADTQEED
GVAKGQVEADSEICVAHTGSQEELAEPDAVGSQTPIASAEETEVGEASDREGIAEAKATVCADAVDACPDQV
EAFEIEKVEDSILDELQTELNAPADKTYEDVLAFDAVCSEALSAFYAVPSDETHFKVCGFYSPAIERTNCWLRST
LIVMQSLPLEFKDLEMQKLWLSYKAGYDQCFVDKLVKSVPKSIILPQGGYVADFAYFFLSQCSFKAYANWRCL
ECDMELKLQGLDAMFFYGDVVSHMCKCGNSMTLLSADIPYTLHFGVRDDKFCAFYTPRKVFRAACAVDVN
DCHSMAVVEGKQIDGKVVTKFIGDKFDFMVGYGMTFSMSPFELAQLYGSCITPNVCFVKGDVIKVVRLVNA
EVIVNPANGRMAHGAGVAGAIAEKAGSAFIKETSDMVKAQGVCQVGECYESAGGKLCKKVLNIVGPDARG
HGKQCYSLLERAYQHINKCDNVVTTLISAGIFSVPTDVSLTYLLGVVTKNVILVSNNQDDFDVIEKCQVTSVAG
TKALSLQLAKNLCRDVKFVTNACSSLFSESCFVSSYDVLQEVEALRHDIQLDDDARVFVQANMDCLPTDWRL
VNKFDSVDGVRTIKYFECPGGIFVSSQGKKFGYVQNGSFKEASVSQIRALLANKVDVLCTVDGVNFRSCCVAE
GEVFGKTLGSVFCDGINVTKVRCSAIYKGKVFFQYSDLSEADLVAVKDAFGFDEPQLLKYYTMLGMCKWPVV
VCGNYFAFKQSNNNCYINVACLMLQHLSLKFPKWQWQEAWNEFRSGKPLRFVSLVLAKGSFKFNEPSDSID
FMRVVLREADLSGATCNLEFVCKCGVKQEQRKGVDAVMHFGTLDKGDLVRGYNIACTCGSKLVHCTQFNV
PFLICSNTPEGRKLPDDVVAANIFTGGSVGHYTHVKCKPKYQLYDACNVNKVSEAKGNFTDCLYLKNLKQTFS
SVLTTFYLDDVKCVEYKPDLSQYYCESGKYYTKPIIKAQFRTFEKVDGVYTNFKLVGHSIAEKLNAKLGFDCNSP
FVEYKITEWPTATGDVVLASDDLYVSRYSSGCITFGKPVVWLGHEEASLKSLTYFNRPSVVCENKFNVLPVDVS
EPTDKGPVPAAVLVTGVPGADASAGAGIAKEQKACASASVEDQVVTEVRQEPSVSAADVKEVKLNGVKKPV
KVEGSVVVNDPTSETKVVKSLSIVDVYDMFLTGCKYVVWTANELSRLVNSPTVREYVKWGMGKIVTPAKLLL
LRDEKQEFVAPKVVKAKAIACYCAVKWFLLYCFSWIKFNTDNKVIYTTEVASKLTFKLCCLAFKNALQTFNWS
VVSRGFFLVATVFLLWFNFLYANVILSDFYLPNIGPLPTFVGQIVAWFKTTFGVSTICDFYQVTDLGYRSSFCNG
SMVCELCFSGFDMLDNYDAINVVQHVVDRRLSFDYISLFKLVVELVIGYSLYTVCFYPLFVLIGMQLLTTWLPE
FFMLETMHWSARLFVFVANMLPAFTLLRFYIVVTAMYKVYCLCRHVMYGCSKPGCLFCYKRNRSVRVKCST
VVGGSLRYYDVMANGGTGFCKHQWNCLNCNSWKPGNTFITHEAAADLSKELKRPVNPTDSAYYSVTEVKQ

VGCSMRLFYERDGQRVYDDVNASLFVDMNGLLHSKVKGVPETHVVVVENEADKAGFLGAAVFYAQSLYRPMLMVEKKLITTANTGLSVSRTMFDLYVDSLLNVLDVDRKSLTSFVNAAHNSLKEGVQLEQVMDTFIGCARRKCAIDSDVETKSITKSVMSAVNAGVDFTDESCNNLVPTYVKSDTIVAADLGVLIQNNAKHVQANVAKAANVACIWSVDAFNQLSADLQHRLRKACSKTGLKIKLTYNKQEANVPILTTPFSLKGG

Constructs: full-length

### Supplementary Figures

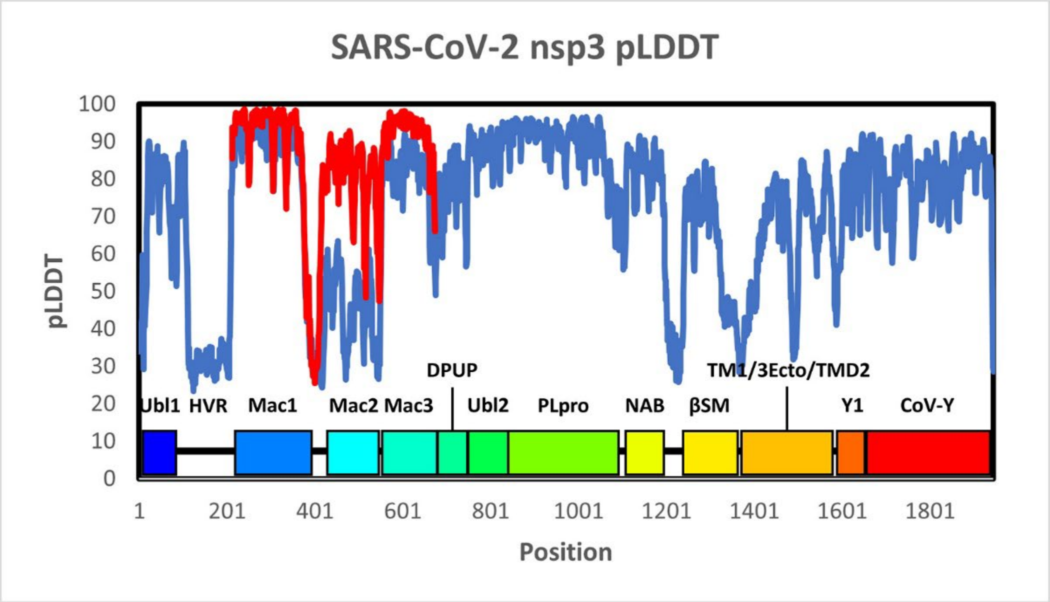


**Figure S1.** AlphaFold per residue confidence for nsp3 as reflected in the predicted Local Distance Difference Test (pLDDT) metric. Scores higher than 70 are generally considered well-predicted. For the full-length construct (shown in blue), most of the protein is well-predicted, with the notable exceptions of the regions spanning residues 111-208 (highly variable region - HVR), 380-550 (Mac2), 1200-1240 (NAB-βSM linker), 1327-1406 (βSM-TM1 linker), and a few shorter linkers between domains. Subsequent models generated for just the Mac1-Mac2-Mac3 region (residues 212-675) dramatically improved the prediction for the Mac2 domain (shown in red).

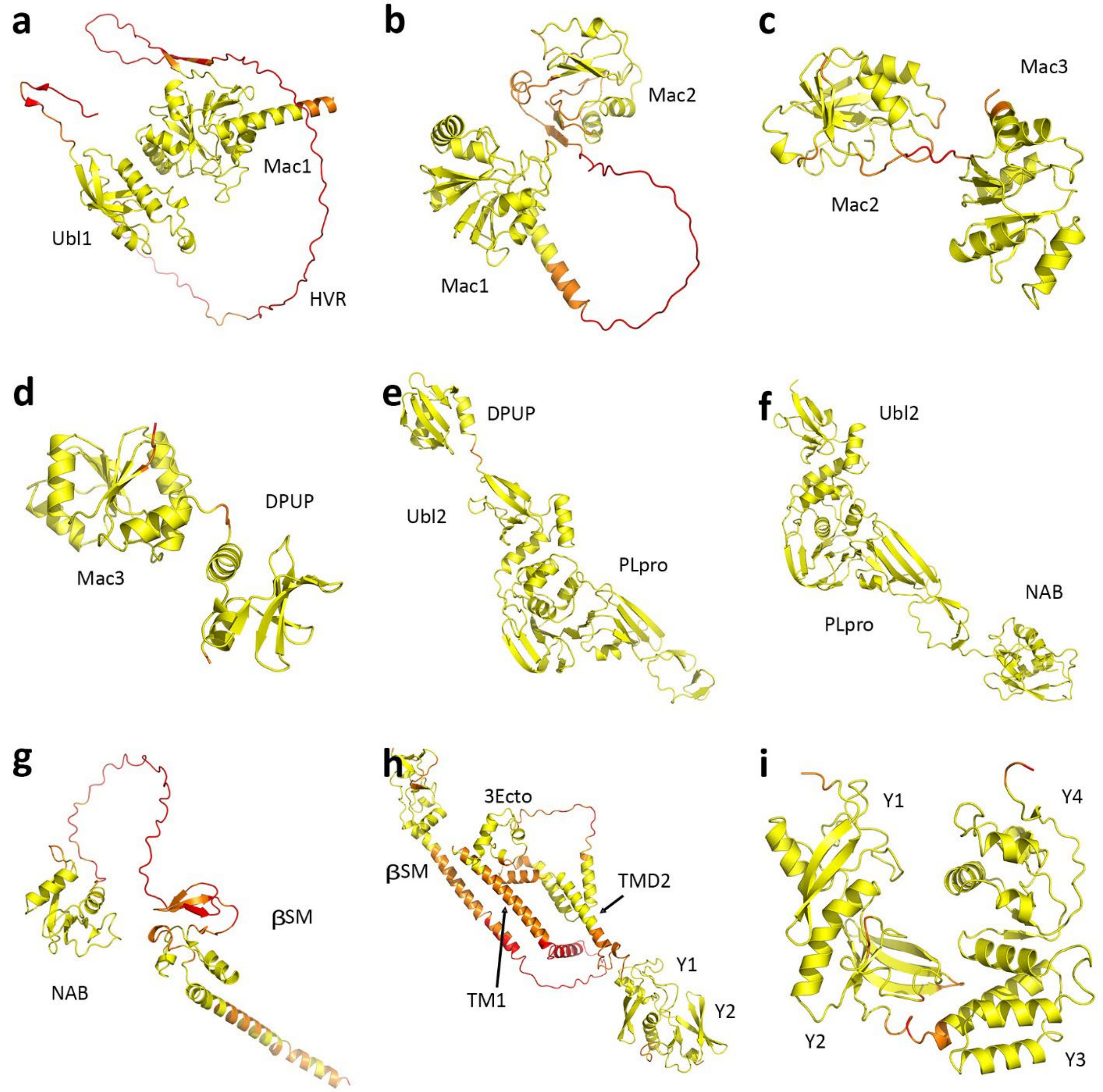


**Figure S2.** Detail of Alphafold predictions of the nsp3 domains. Predictions are not from the full protein but for the specific constructs shown. Each residue is color coded by its pLDDT score (75-100 yellow; 50-75 orange; 0-50 red). The low pLDDT score associated with most of the linkers between domains reflects flexibility in how the domains are oriented with respect to each other. a) Residues 1-387, covering Ubl1, HVR and Mac1, average pLDDT = 77.8. The Ubl1 and Mac1 domains are connected by residues 106-213, which includes the completely unstructured HVR domain. b) Residues 212-539, covering Mac1 and Mac2, average pLDDT = 80.9. The domains are connected by residues 386-416. The Mac2 domain is correctly predicted in this construct. c) Residues 416-674, covering Mac2 and Mac3, average pLDDT = 86.9. The domains are connected by residues 539-549. The Mac2 domain is also correctly predicted in this construct. d) Residues 547-744, covering Mac3 and DPUP, average pLDDT = 90.4. The domains are connected by residues 673-680. e) Residues 678-1093, covering DPUP and Ubl2/PLpro, average pLDDT = 95.1. The DPUP and Ubl2 domains are connected by residues 744-748. f) Residues 747-1196, covering Ubl2/PLpro and NAB, average pLDDT = 95.8. The PLpro and NAB domains are connected by residues 1090-1095. g) Residues 1109-1365, covering NAB and βSM, average pLDDT = 71.2. The two domains are connected by residues 1194-1242. h) Residues 1241-1761, covering βSM, TM1, 3Ecto, TMD2. Y1 and Y2 of the CoV-Y domain, average pLDDT = 73.8. The βSM amphipathic helix and TM1 do not form any tight bundle, as reflected in the lower pLDDT scores. i) Residues 1596-1945, covering Y1 and the CoV-Y domain (Y2, Y3 and Y4), average pLDDT = 89.8. Y2 and Y3 are connected by residues 1756-1764, which introduces the most flexibility in the domain.

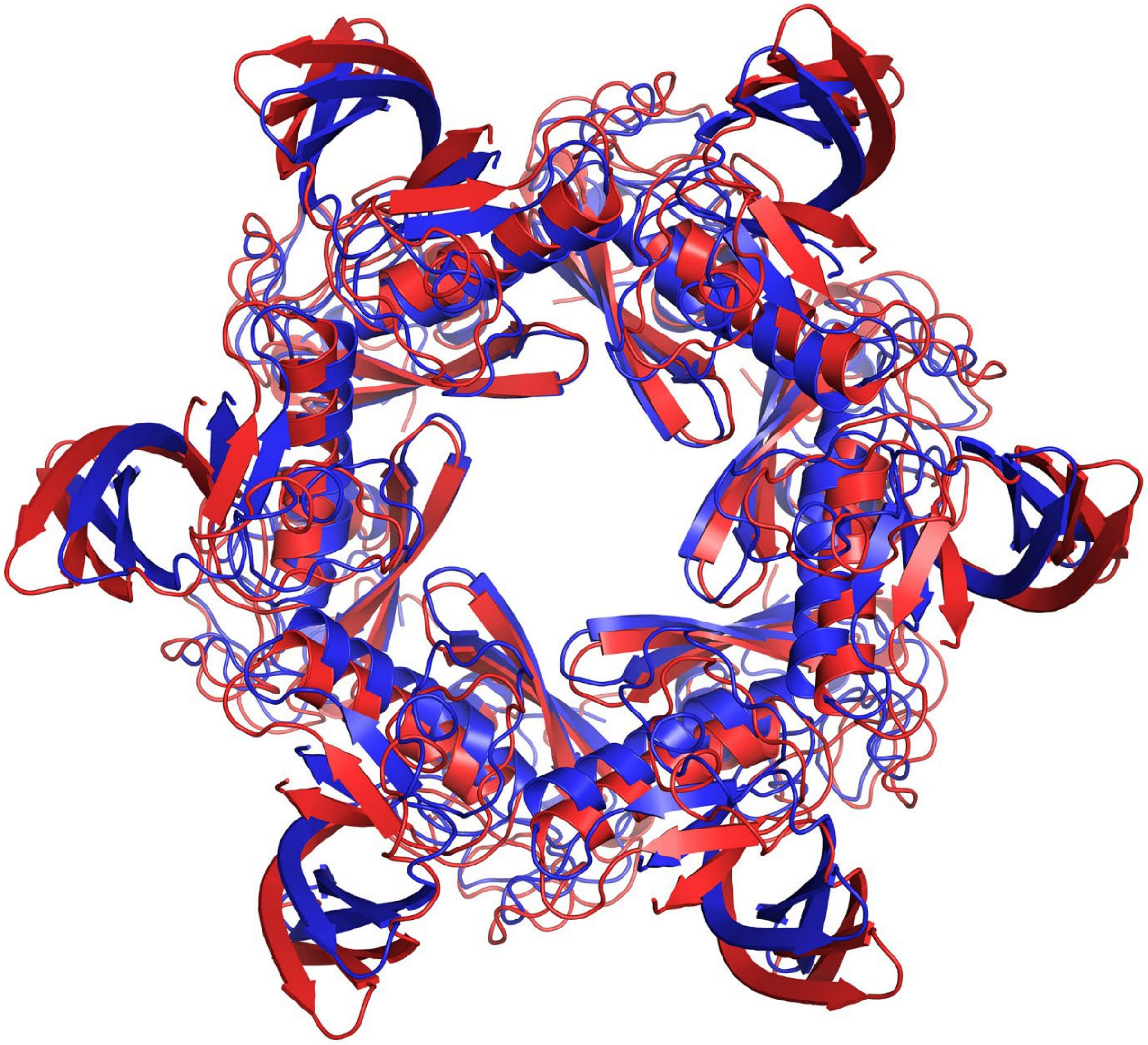

**Figure S3.** Overlay of AlphaFold prediction of hexameric nsp3 Y1/Y2 (residues 1587-1756, shown in blue) with the central nsp3 pore component from the cryo-ET structure of Huang, et al. (shown in red).(1) Multiple hexameric constructs which included the Y1/Y2 domains (including the full CTD, residues 1587-1945) generated this same structure.

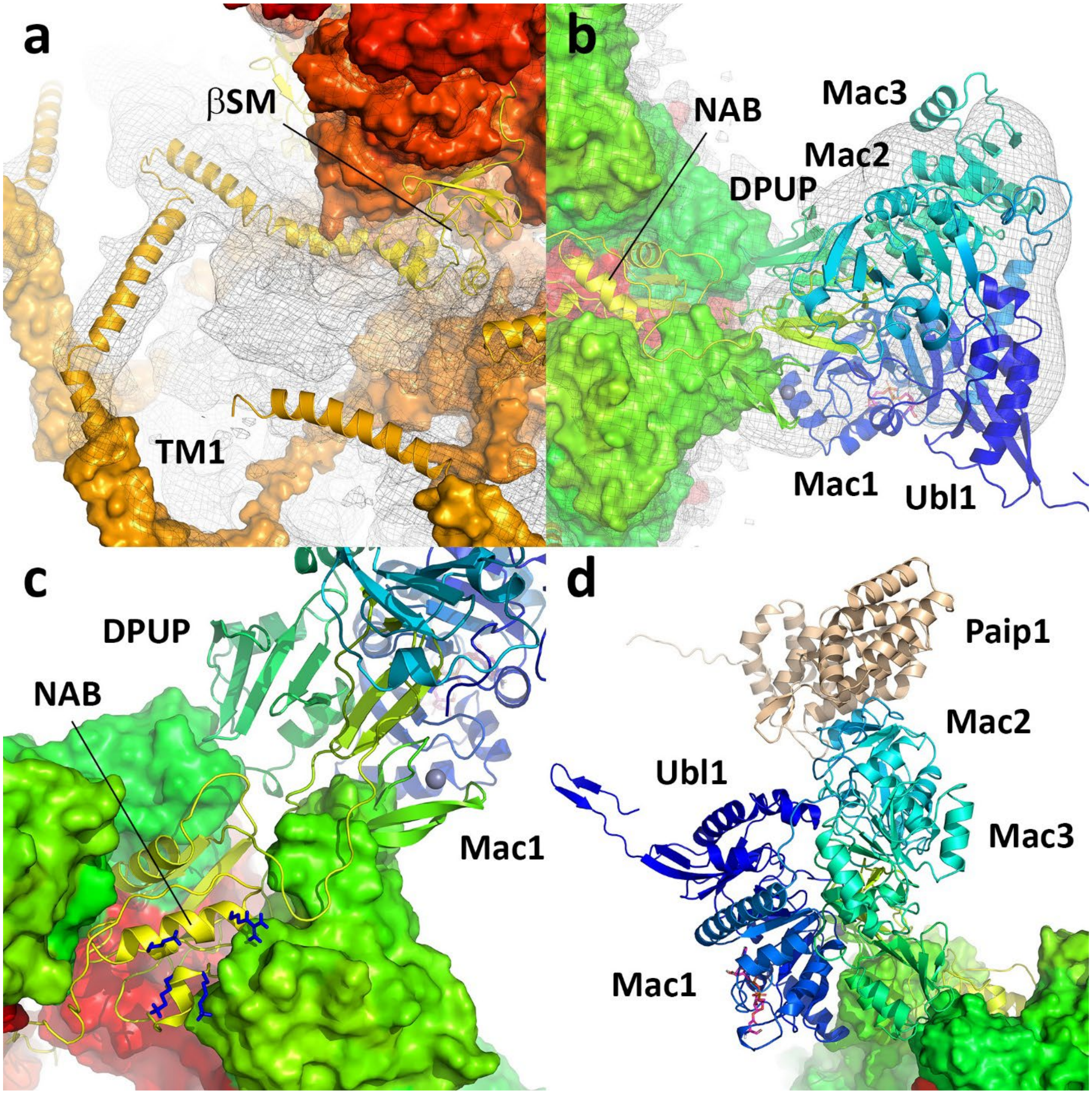


**Figure S4.** Detail of the remodeled nsp3 component of the pore. a) Alternating subunits exhibit unfit density on the surface of the outer membrane consistent with placement of two amphipathic helices which connect the βSM domain to TM1. The βSM domain is placed at the base of the crown, although density is unclear. b) The prongs were remodeled, composed of Ubl1, Mac1, Mac2, Mac3 and DPUP domains. c) The NAB domain was placed in the rim of the crown between PLpro domains. Basic residues which constitute the RNA binding motif may serve to direct nascent RNA toward the prongs. d) The Mac1 domain is positioned at the base of the prong, with its ADPR active site directed outward. A substrate fragment is shown in pink to indicate the active site location. The Mac2 domain sits at the top of the prong and coordinates the translation cofactor Paip1, shown in beige.

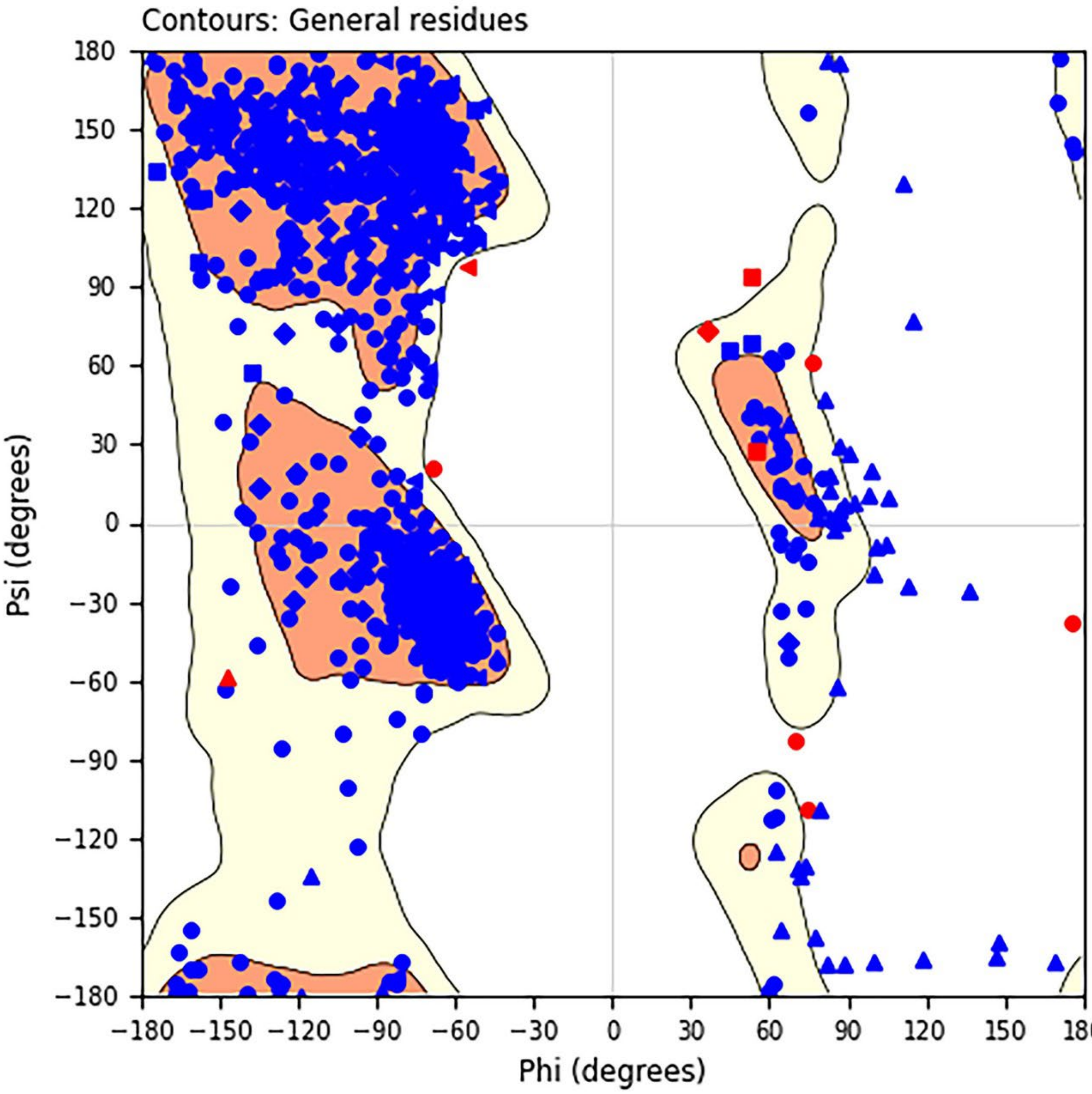


**Figure S5.** Ramachandran analysis of the remodeled nsp3 N-terminal prong (residues 1-111 and 204-1212). Glycine is plotted as triangles, proline as arrows, pre-proline as squares, isoleucine/valine as diamonds, and all other residues as circles. The orange regions are the "favored" regions for general residues and the yellow regions are considered "allowed". 1.43% of phi/psi angles are considered disallowed in this analysis.

352

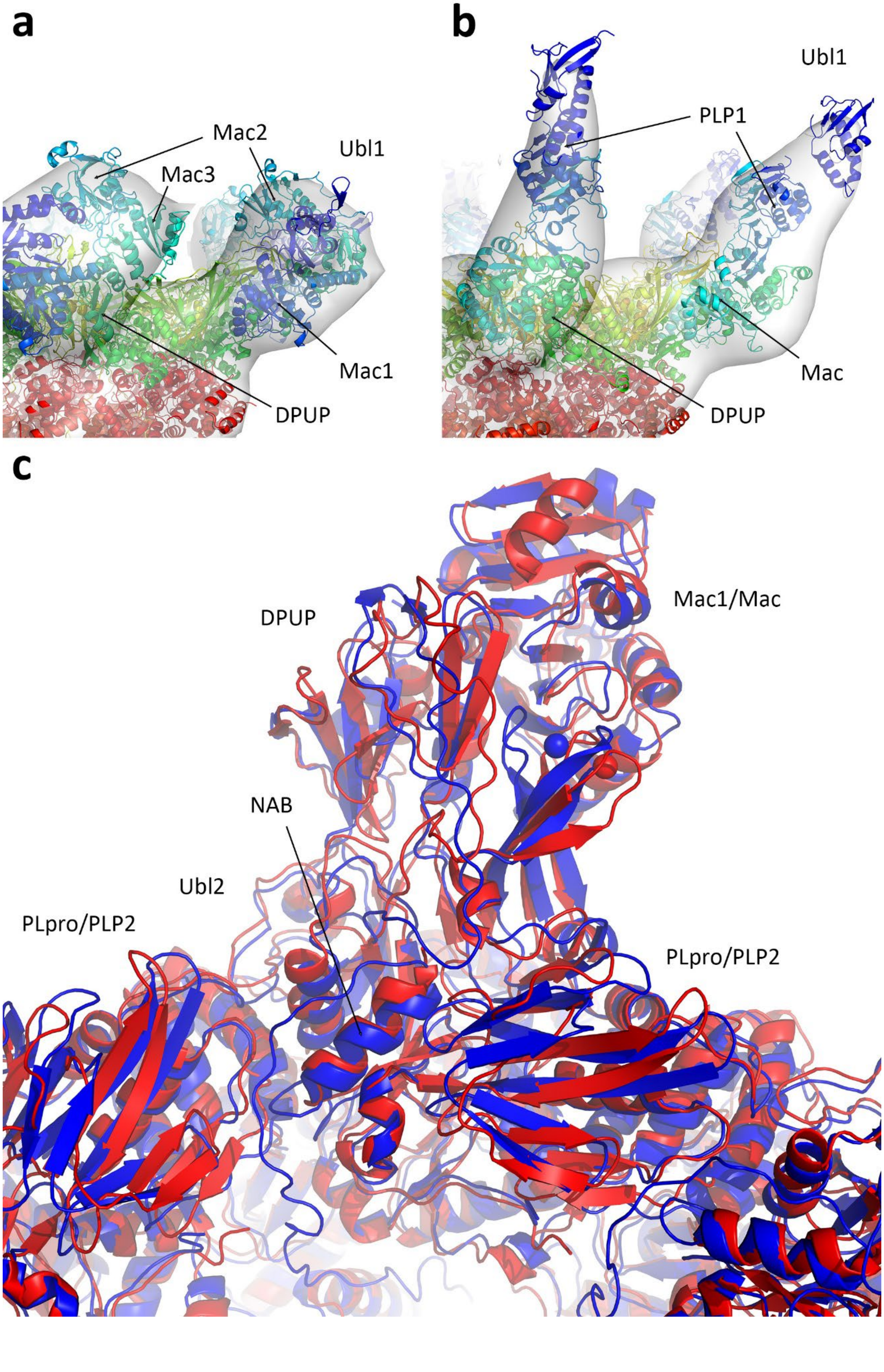
a
Mac2
Mac3
Ubl1
Mac1
DPUP
b
Ubl1
PLP1
Mac
DPUP
c
Mac1/Mac
DPUP
NAB
Ubl2
PLpro/PLP2
PLpro/PLP2

**Figure S6.** Comparison of the models for the dodecameric nsp3 crowns of SARS-CoV-2 and MHV. a) Detail of the remodeled SARS-CoV-2 prong shown against the cryo-ET map from Zimmermann, et al.(2) corrected for handedness (see Supplementary text). b) Detail of the modeled MHV prong shown against the cryo-ET map from Wolff, et al.(3) Differences in the shapes of the prongs stem from differences in the N-terminal domains. c) Overlay of SARS-CoV-2 (blue) and MHV (red) models highlighting the conserved domains between the two viruses which form the rim of the cytosolic crown and the base of the prongs. The Ubl1, HVR, Mac2 and Mac3 domains of SARS-CoV-2 and the Ubl1, HVR and PLP1 domains of MHV are not shown.

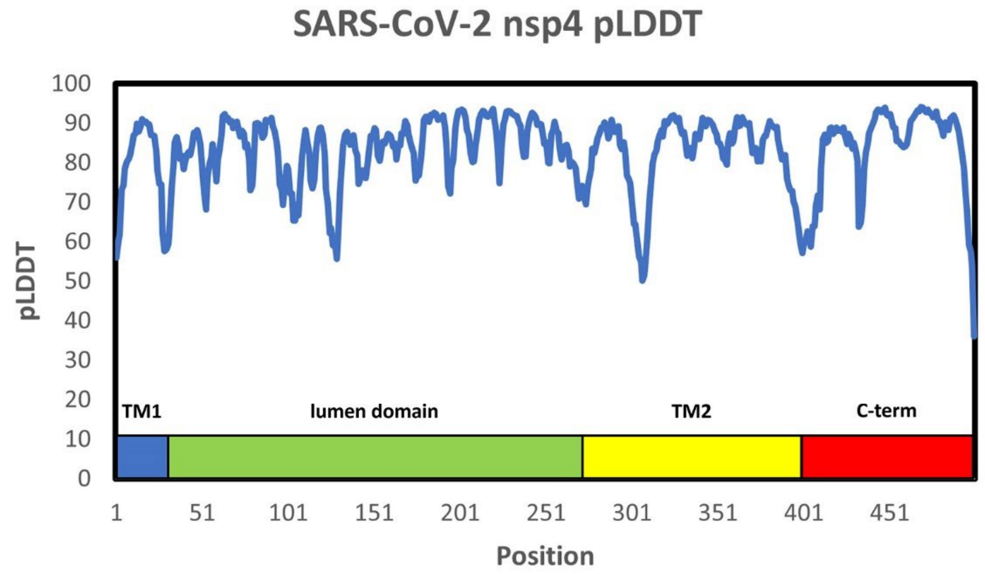


**Figure S7.** AlphaFold per residue pLDDT confidence for nsp4. The protein can be considered well-predicted.

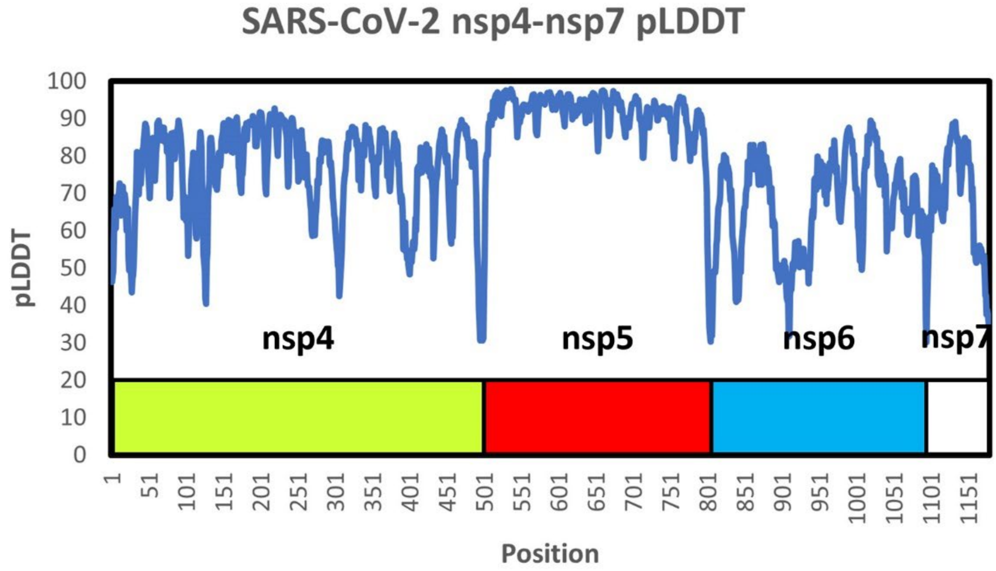


**Figure S8.** AlphaFold per residue pLDDT confidence for nsp4-nsp7. The polyprotein can be considered generally well-predicted.

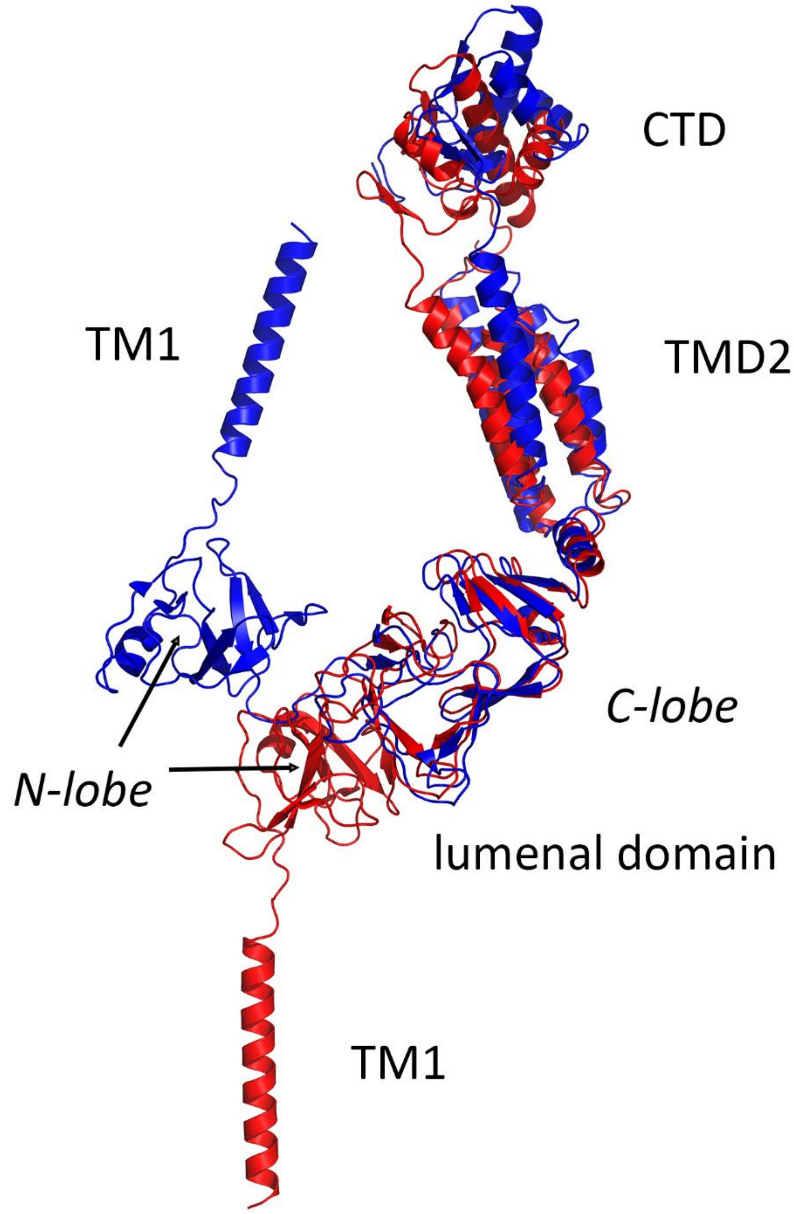


**Figure S9.** Overlay of two conformations of the nsp4 protein. The conformation in blue is consistent with binding to a single membrane and is derived from the Alphafold prediction for the nsp4-nsp5-nsp6-nsp7 uncleaved polyprotein. The conformation in red is consistent with binding to a double membrane and is predicted for the cleaved nsp4 protein. The conformational change is driven by the lumenal domain, where the N-lobe and C-lobe are more loosely associated when bound to a single membrane, but tightly associated when bound to a double membrane.

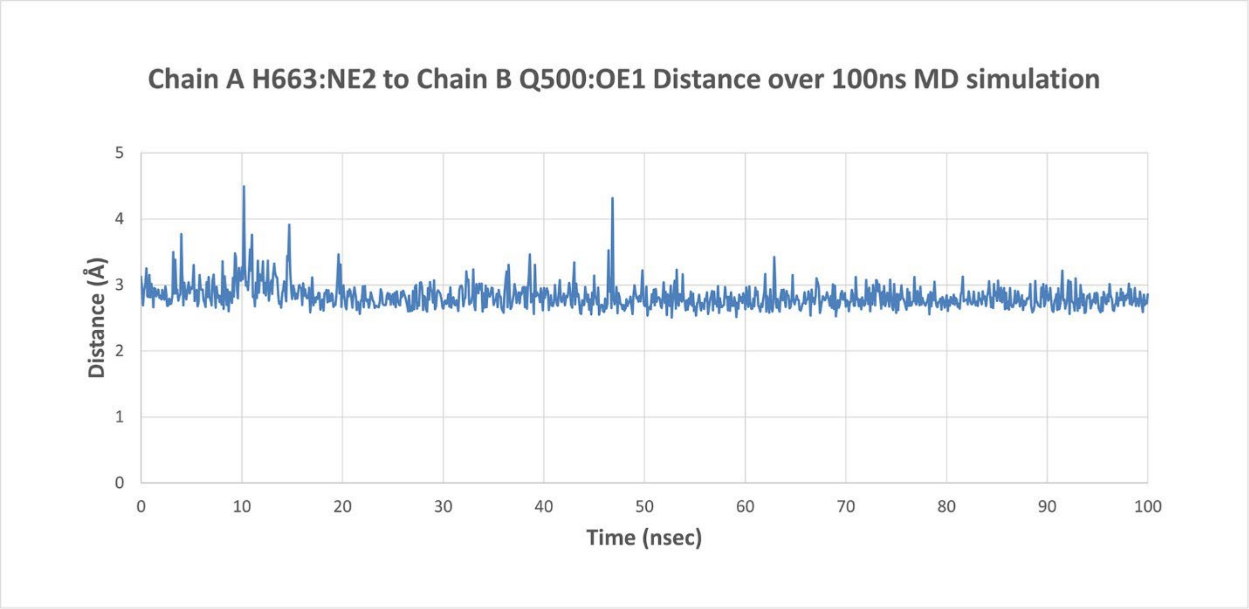


**Figure S10.** Measurement of the distance between the Chain B nsp4 P1 Q500 OE1 atom and the Chain A nsp5 S1 pocket H663 NE2 atom across a 100 ns MD simulation of the nsp4-nsp5-nsp6-nsp7 uncleaved polyprotein dimer. The Chain B nsp4-nsp5 cleavage site residues are not properly engaged with the active site for cleavage to occur. But the Q500 residue effectively blocks the S1 pocket from engaging any other substrate, rendering the protease inactive.

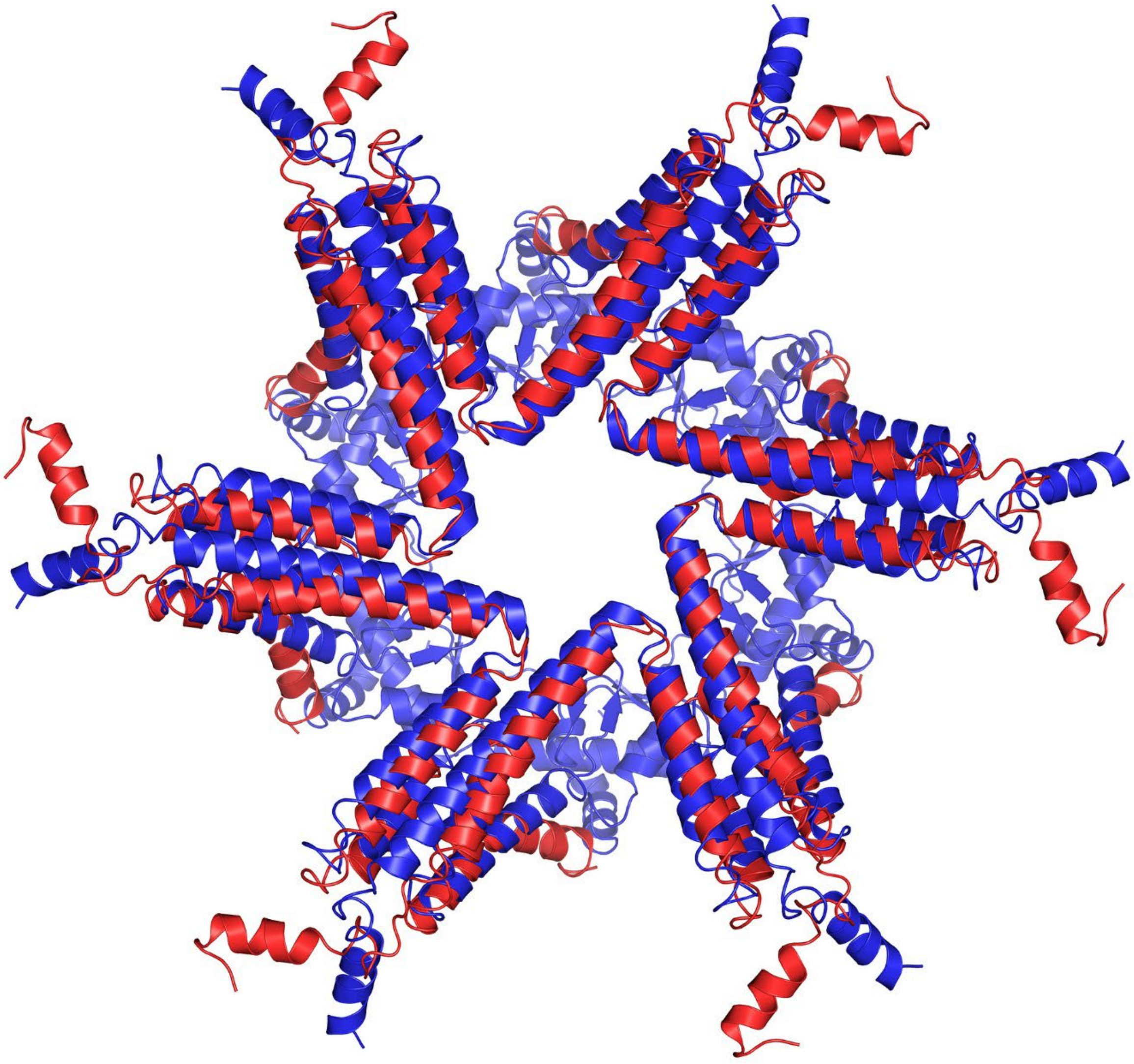

**Figure S11.** An overlay of the central nsp4 hexameric pore component from the cryo-ET structure (PDB 8YAX, red) and that predicted by AlphaFold (blue). The cryo-ET structure is an extraction of residues 256-401 from the inner nsp4 hexamer. The AlphaFold prediction used a construct covering residues 256-500. Note, the outer nsp4 hexamer was not predicted by Alphafold and is not shown for clarity.

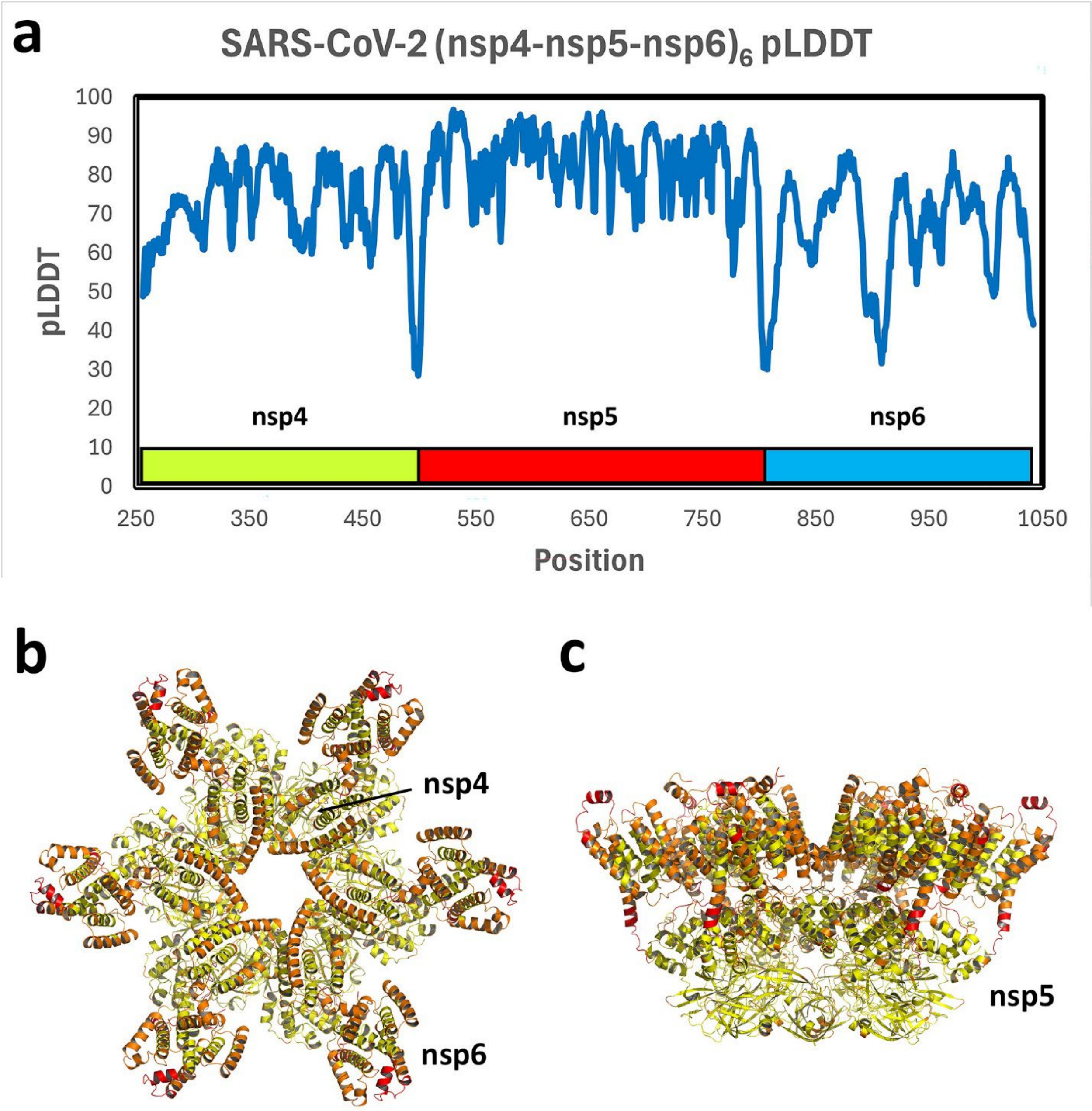


**Figure S12.** a) AlphaFold per residue pLDDT confidence for the predicted nsp4-nsp5-nsp6 hexamer (residues 256-1042). b) Top view of predicted structure color coded by the pLDDT ((75-100 yellow; 50-75 orange; 0-50 red). c) Side view of the predicted structure.

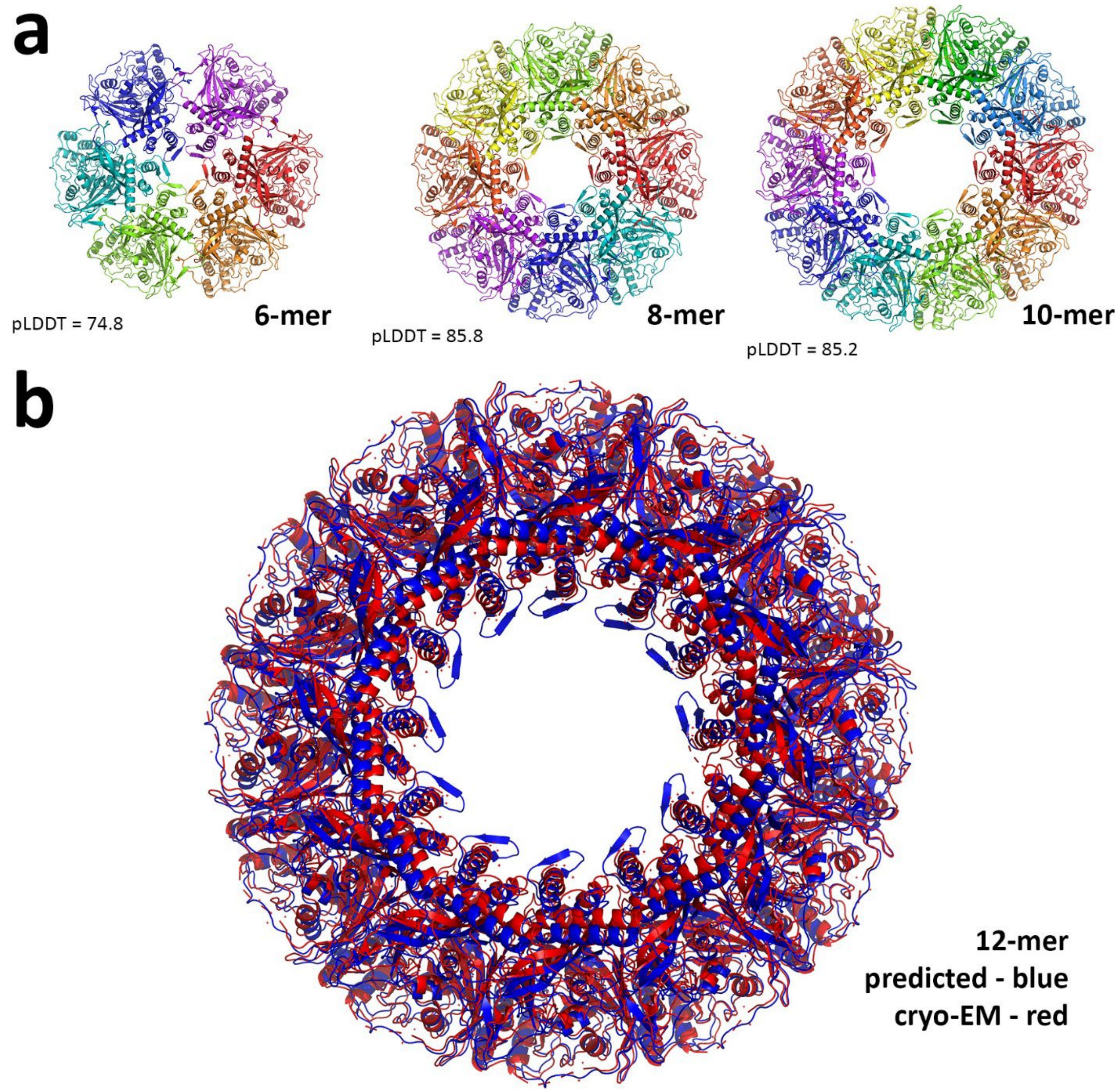


**Figure S13.** a) Alphafold predictions of the hexamer, octamer and dodecamer of chikungunya virus nsP1. The packing in each remains similar, forming rings of increasing size and density. b) Comparison of a predicted nsP1 dodecamer to the cryo-EM structure (6Z0U).(9) The prediction was made based on an extrapolation from the Alphafold hexamer, using a target packing distance determined from the predicted dimer (not shown).

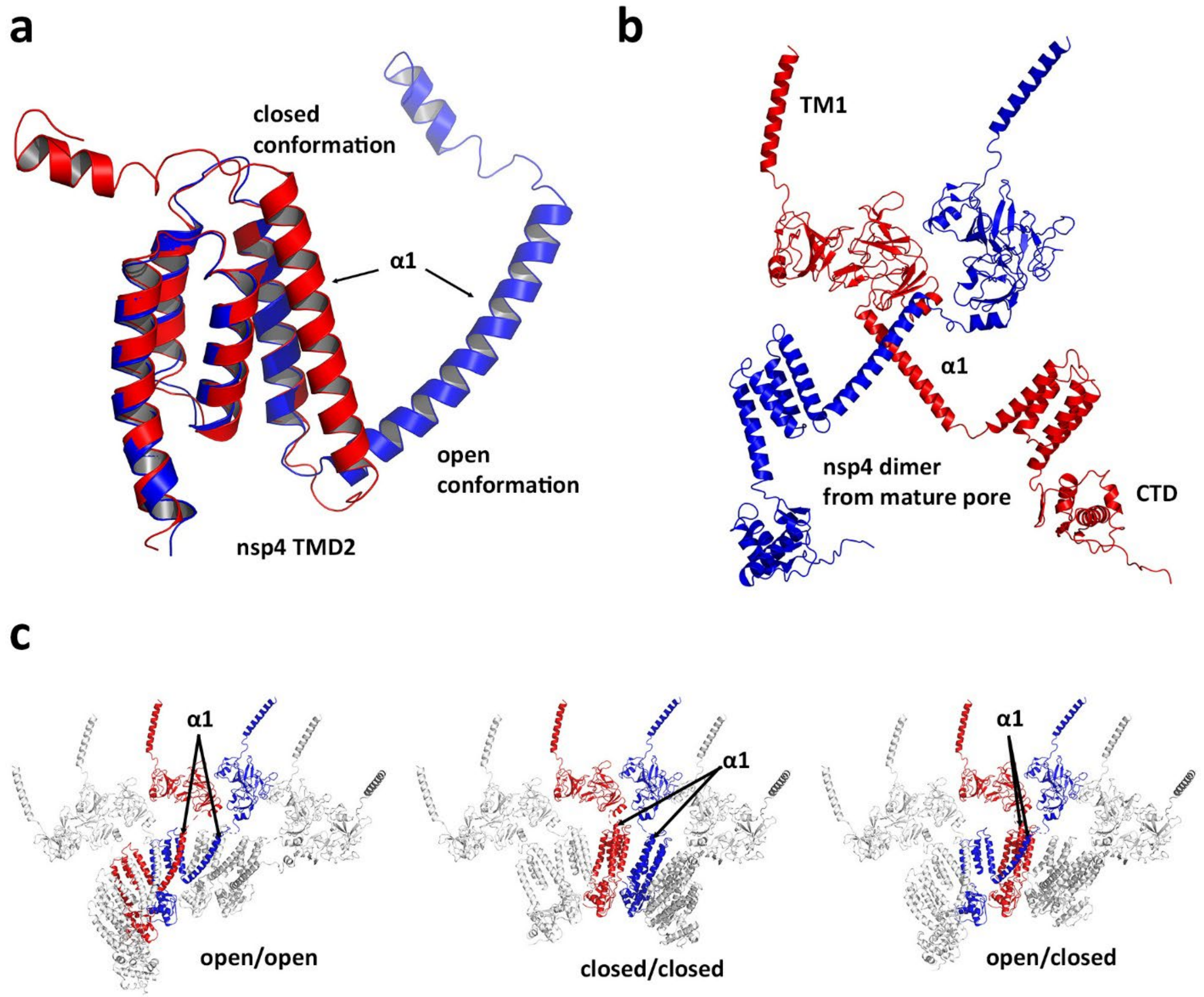


**Figure S14.** a) Comparison of Alphafold predictions for the nsp4 TMD2 (residues 256-399) from the nsp4 monomer (red) and the nsp4-nsp5-nsp6 hexamer (blue). The α1 helix adopts a closed conformation in the monomer and an open conformation in the hexamer. b) An nsp4 dimer extracted from the cryo-ET pore structure of Huang, et al.(1) The subunit in blue is part of the inner nsp4 hexamer, while that in red is part of the outer nsp4 hexamer. Both adopt an open conformation, but the subunit in blue must cross in front of the one in red. c) Potential conformations of nsp4 within the proposed dodecamer uncleaved polyprotein. The structures differ only with respect to the conformation of the α1 helix. The mixed open/closed arrangement has the most direct route to evolve to the final pore structure upon nsp4/nsp5 cleavage.

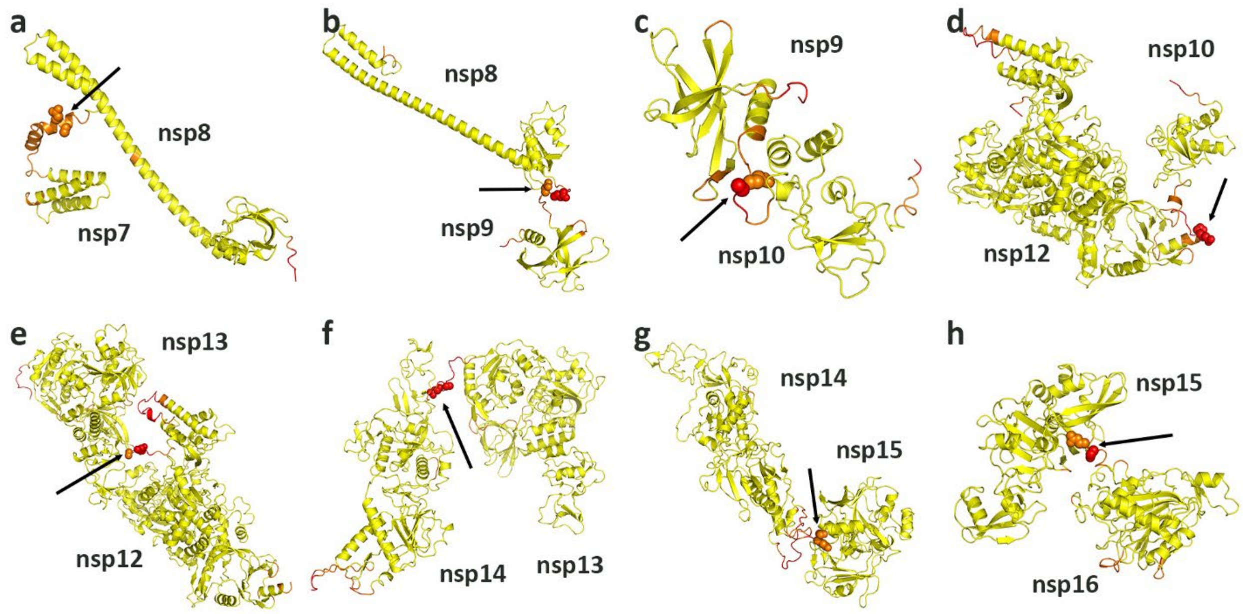


**Figure S15.** Alphafold predcitions of the uncleaved polyprotein. Predictions are for the specific constructs shown. Each residue is color coded by its pLDDT score (75-100 yellow; 50-75 orange; 0-50 red), which highlights the flexibility of the linker between each nsp protein pair. a) nsp7-nsp8 (residues 1097-1377, average pLDDT = 87.5). The linker spans residues 1157-1190 and is largely helical in character but conformationally flexible. b) nsp8-nsp9 (residues 1180-1490, average pLDDT = 89.2). The linker spans residues 1370-1386 and is an unstructured loop. c) nsp9-nsp10 (residues 1378-1629, average pLDDT = 85.5). The linker spans residues 1487-1500 and is an unstructured loop. d) nsp10-nsp12 (residues 1491-2561, average pLDDT = 92.2). The linker spans residues 1623-1646 and has some helical character. e) nsp12-nsp13 (residues 1630-3162, average pLDDT = 90.1). The linker spans residues 2529-2563 and has some helical character. f) nsp13-nsp14 (residues 2562-3689, average pLDDT = 89.6). The linker spans residues 3151-3171 and is an unstructured loop. g) nsp14-nsp15 (residues 3163-4035, average pLDDT = 91.9). The linker spans residues 3683-3691 and is an unstructured loop. h) nsp15-nsp16 (residues 3690-4333, average pLDDT = 92.5). The linker spans residues 4032-4043 and is an unstructured loop.

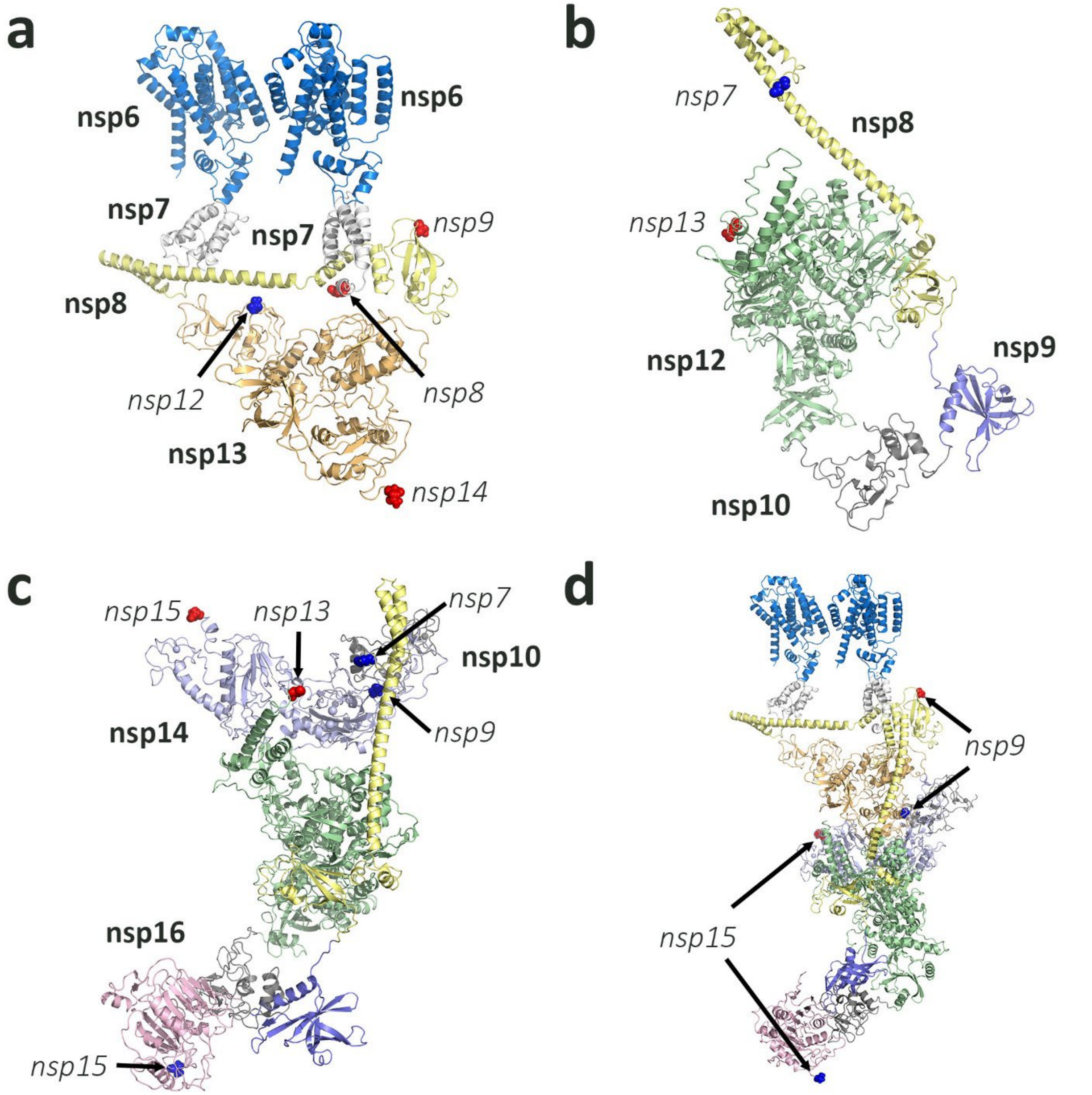


**Figure S16.** The pp1a'/pp1ab' uncleaved polyprotein dimers were constructed from existing complex structures of the cleaved proteins and Alphafold models. a) The predicted nsp4-nsp5-nsp6 dodecamer was used as an anchor. It was first extended to include nsp7, based on the Alphafold prediction for the nsp4-nsp5-nsp6-nsp7 monomer. In the extracted dimer shown, one subunit could be further extended to include nsp8 and this could interact with the second nsp7 and recruit nsp13 just as seen in the cryo-EM structures of the nsp12/nsp7/$(nsp8)_2$/$(nsp13)_2$ complex (6XEZ). The locations of the linkage points for additional proteins are indicated in italics. b) An Alphafold prediction of uncleaved nsp8-nsp9-nsp10-nsp12 showed nsp8 interacts with nsp12 as in the replication complex (6XEZ). The positioning of nsp9 and nsp10 is highly flexible. c) nsp14/nsp10 was coordinated to the nsp8-nsp9-nsp10-nsp12 polyprotein following previous predictions for the full hexameric replication complex.

nsp16 coordinates to nsp10 as seen in multiple x-ray structures (6W4H). d) The models from a) and c) were linked leaving only nsp8 to be connected to nsp10 with nsp9 and nsp14 to be connected to nsp16 with nsp15.

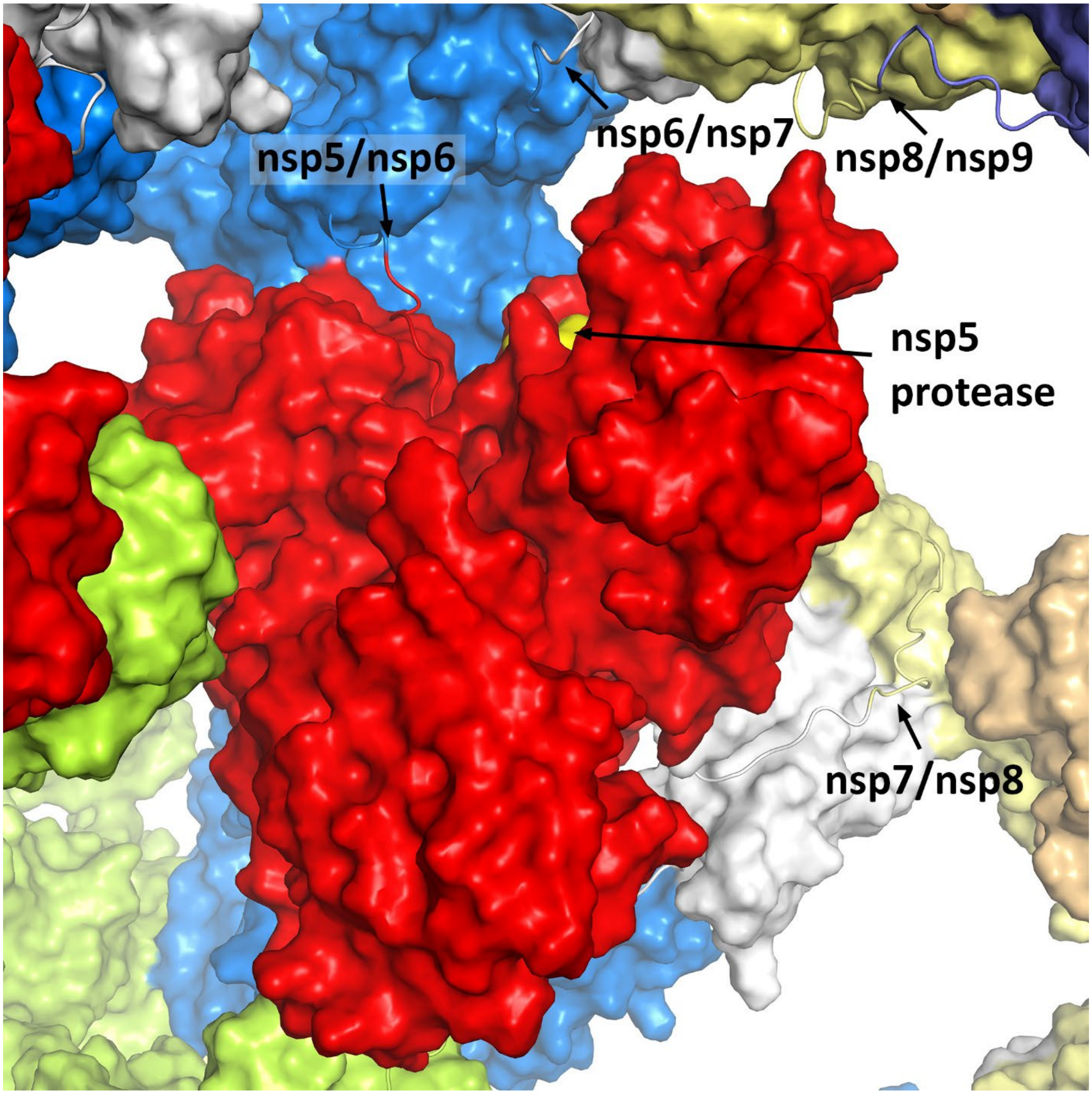


**Figure S17.** Cleavage at alternating nsp4-nsp5 linkers in the dodecamer facilitates pore maturation and leads to formation of canonical nsp5 dimers. The protease active site of the uncleaved subunit is activated by the dimerization, and several additional cleavage sites are within range.
